# The scales and signatures of climate adaptation by the *Arabidopsis* transcriptome

**DOI:** 10.1101/2020.10.28.358325

**Authors:** Jack M. Colicchio, Melis Akman, Benjamin K. Blackman

## Abstract

Clines in allele frequency and trait variation can be highly informative for understanding how populations have historically adapted to climate variation across landscapes. However, as a consequence of the many complexities inherent to this process, these climate-associated differentiation patterns can be confounded, misleading, or obscured. Molecular phenotypes like gene expression levels are a potentially valuable means for resolving these complexities. Their intermediate position between genomes and organismal traits and their interrelatedness structured by gene regulatory networks can help parse how different climatic factors contribute to unique components of range-wide or region-specific diversity patterns. Here, we demonstrate these explanatory values of gene expression variation through integrative analyses of transcriptomic data from 665 *Arabidopsis thaliana* accessions. Differentiation of co-expressed genes is often associated with source site climate. Although some patterns hold range-wide, many other gene expression clines are specific to particular ancestry groups, reflecting how broad-scale and local combinations of selective agents differentially resolve functional interrelationships between plant defense, drought tolerance, and life history traits. We also extend these analyses to parse how different factors explain climate-associated variation in flowering time and its plasticity. Expression of key regulators *FLC* and *SOC1* strongly predicts time to flower, consistent with previous work, but our results also highlight novel relationships that indicate as yet unexplored climate-related connections between defense signaling and flowering. Finally, we show that integrative models combining genotype and gene expression information predict variation in flowering time under ecologically realistic conditions more accurately than models based on either source alone.

**Significance Statement:** Populations often adapt to local conditions along climate gradients, and associations between climate parameters and traits or alleles often indicate a history of adaptive differentiation. However, such signals can be obscured or misleading due to the complex genetics underlying trait variation or other historical processes, frustrating our capacity to reveal how populations adapt to diverse climates. As a molecular intermediate between genetic polymorphisms and their impact on organismal traits, gene expression variation is a useful readout for addressing several of these difficulties. Here, we leverage transcriptomic data from hundreds of *Arabidopsis thaliana* accessions to reveal continental and region-specific patterns of climate-associated differentiation as well as investigate how gene expression adaptation at both scales shapes flowering time variation along climate gradients.

## Introduction

Determining what combinations of traits and genetic variants adapt populations to their local climates is an essential pursuit in evolutionary biology, as knowledge of how populations have coped with past or recent environmental challenges improves our capacity to predict whether and how populations will evolve to persist in future climates. Since early studies of clinal variation (1–4), many approaches that link variation in the climate parameters that act as agents of selection with locally adaptive phenotypic and genetic variation have highlighted the complex nature of this evolutionary process. Local adaptation is often polygenic (5, 6), may involve the evolution of covarying traits (or trait syndromes) (7), and may result in different populations exploiting distinct ecological strategies to reach similar local fitness peaks (8). These complexities pose many analytical challenges (9, 10). For instance, the polygenic nature of trait variation necessitates large sample sizes to detect minor effect or rare alleles in association mapping studies. Covariation among allele frequencies (linkage disequilibrium), among phenotypes that comprise trait syndromes, among environmental factors, and among all these factors and population structure often stymies our abilities to make specific inferences about which causal forces have spurred ecological diversification at the phenotypic or genotypic level and to construct genotype-to-phenotype maps. In addition, regional adaptive signatures can obfuscate adaptive patterns on a global scale. Several creative solutions to these challenges have been developed, including restricting focus to population samples with limited population structure (11, 12), distilling multidimensional relationships among traits or climatic parameter to primary axes of variation (13, 14), and sampling across replicate environmental gradients to assess evolutionary convergence (8, 15).

Here, we further surmount these obstacles through analyzing natural variation in gene expression in tandem with climatic, genotypic, and phenotypic data all available for a large set of *Arabidopsis thaliana* accessions (16). Gene expression levels are intermediate molecular phenotypes, and changes in gene expression have often been highlighted as a common mechanism for adaptive evolution (17–20). As for any complex trait, gene expression variation among individuals or populations may reflect genetic polymorphism, developmental differences, environmental variation, or interactions among these factors (17, 18, 21–23). The hierarchical and integrative structure of gene regulatory networks confers several advantages that can facilitate connecting polygenic variation to phenotype and environment. For instance, allele-specific expression and expression quantitative trait locus (eQTL) studies have used evolutionary sign tests to identify suites of functionally related genes that share consistent patterns of evolutionary change between strains, populations, or species (17, 24–26).

Likewise, gene expression is a helpful intermediate for studying local adaptation along environmental gradients even when this process has involved minor shifts in allele frequencies across many polymorphisms in single or multiple loci. Such allelic variation can be especially challenging to associate with phenotype or environment when different polymorphisms have substitutable impacts on phenotype and fitness, lead to parallel adaptation (27, 28), or co-vary with population structure (29). Together, these variants may nonetheless accrete to yield strong environmental clines in gene expression or gene expression plasticity at the immediately affected genes or downstream integrators of their function (19, 21, 30–32). Thus, another effective method for more deeply examining signals of local adaptation is by interrogating environmental associations among genes with co-varying expression levels clustered into gene co-expression networks (33). With sufficient data, gene co-expression modules can then be connected to environmental agents of selection through clinal analysis (34–37), to underlying polymorphisms through association mapping (38), and to organismal phenotypes through correlational tests or analyses of enrichment for particular gene functions (39, 40). Moreover, different co-expression modules may orthogonally influence variation in subcomponents of a more integrated trait or trait syndrome. Taking such a multi-level approach may reveal when adaptation to the same environmental factor has involved different underlying mechanisms at different geographic scales.

Here, we implement a three-tiered framework to test for association between climate and gene expression while controlling for patterns of relatedness (Fig. 1). Our approach detects not only patterns consistent with climate adaptation that have emerged on a continental scale but also region-specific patterns that are often missed or limit the power of standard association approaches. Notably, the signatures of adaptation detected at these different scales are often non-overlapping and distinguished by contrasting impacts on variation in genes at different topological levels within co-expression networks. In addition, building upon ample previous work (41, 42), we describe how known and novel transcriptomic components of flowering time variation relate to underlying genetic variants, and build structural equation models to assess how these various molecular components of flowering time variation are integrated to yield widespread and lineage-specific patterns of climate adaptation (Fig. 1). By linking differentiation in gene expression patterns with associated climatic and trait variation, our findings demonstrate the value of broadly sampled gene expression datasets for dissecting how selection by multifarious environmental agents acts to shape gene regulatory networks and yield adaptive variation in ecological strategies (7).

**Figure 1.**
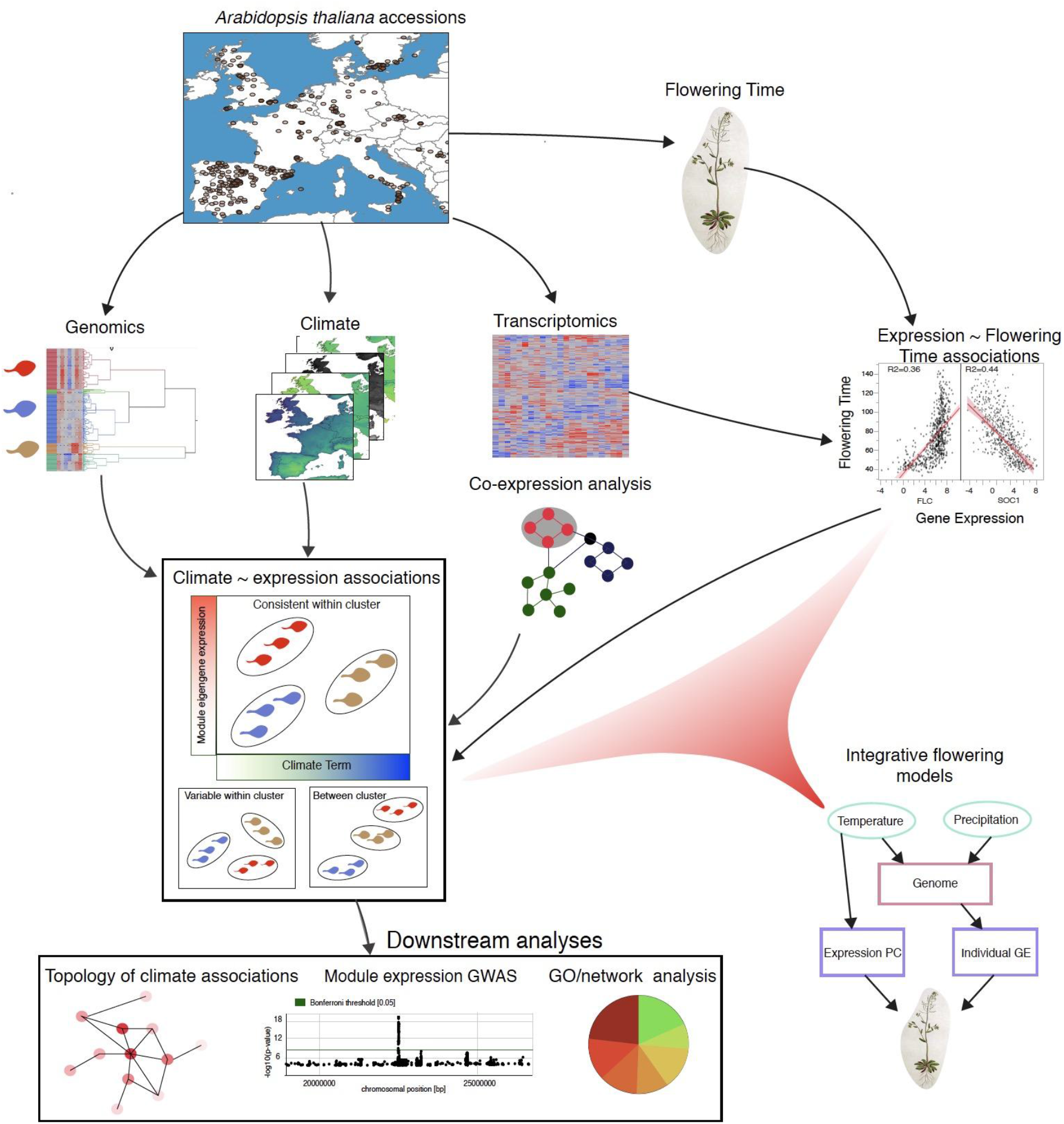
Schematic illustrating integrative approach applied to identify signatures of climate adaptation by the *Arabidopsis* transcriptome and relate patterns of SNP and gene expression variation to flowering time variation.

## Results and Discussion

### Genomic cluster assignment

To account for relatedness in downstream analyses, we clustered *Arabidopsis thaliana* accessions into thirteen groups ranging in size from 8 to 109 accessions using whole genome SNP data (43) (Fig. 2a). A post-hoc discriminant analysis based on the PC values correctly classified 96.3% of ecotypes to their cluster, and the majority of accessions in each of the 13 genomic clusters were assigned to a single admixture group of the set previously defined by the 1001 Genomes project, confirming our approach’s robustness (Fig. S1, Table S1; (43)). Genomic differentiation patterns were largely explained by geography (Fig. 2b); a model including only latitude and longitude correctly classified 77.1% of the accessions to genomic clusters. Using a predictor screening approach, we determined that including temperature seasonal sine deviance (SSD, see *Methods),* temperature annual range, summer precipitation, and winter precipitation all significantly improved the classification rate (82.2%, Table S2).

**Figure 2.**
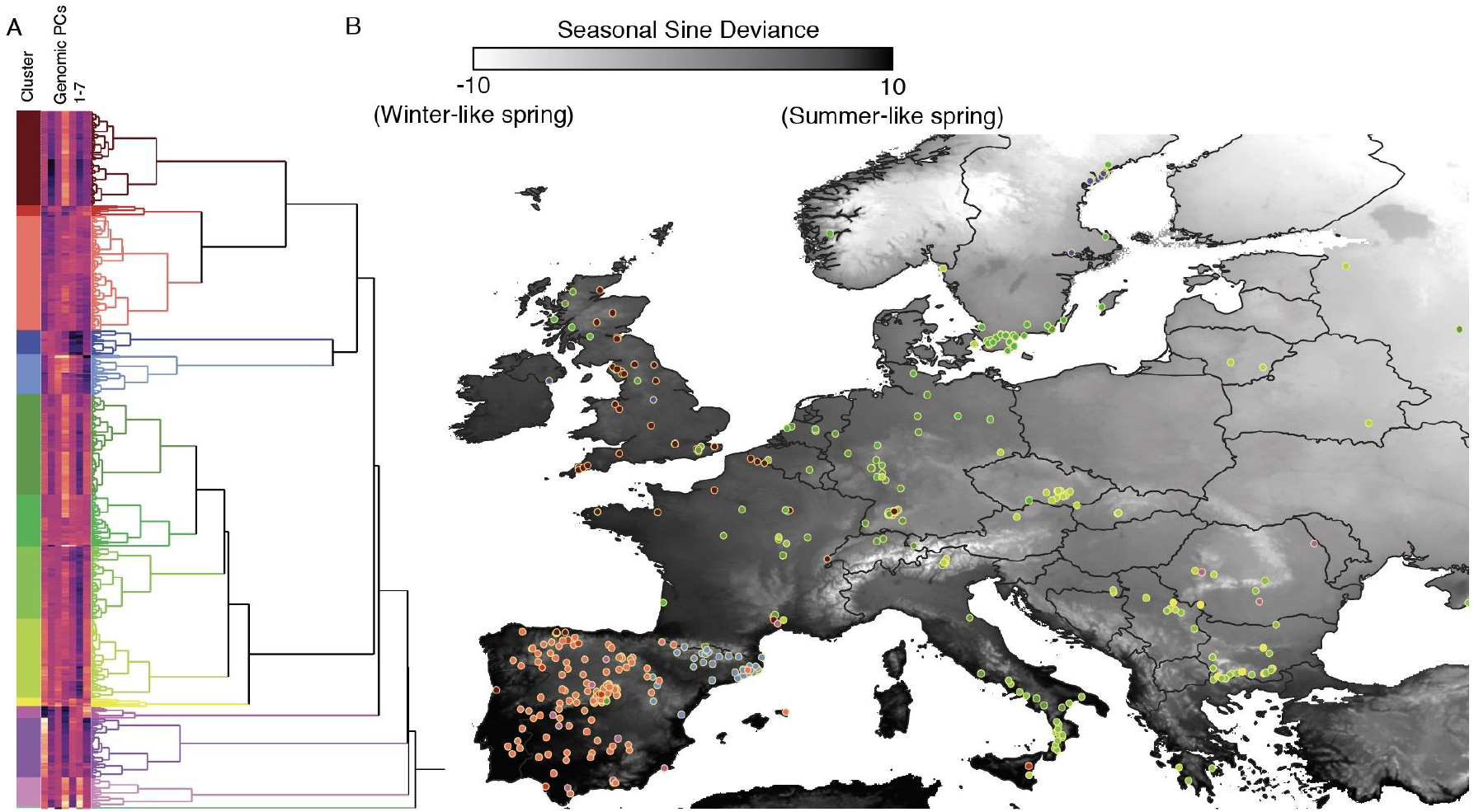
Genomic PC based clustering of Eurasian *Arabidopsis* accessions. (A) Ward clustering on the first 7 PCs of genomic variation identified 13 genomic clusters, largely overlapping with the admixture groups constructed as part of the 1,001 genomes project (Fig. S1). (B) When accessions (colored by cluster membership) are plotted across the landscape, genomic clusters show strong but not complete geographic differentiation. Map shaded by seasonal-sine deviance (SSD), a metric that indicates the deviance from seasonal temperature fluctuations from a sine curve model of seasonal temperature fluctuations. Positive values indicate “broad shoulders” where spring and fall months are warmer, while negative values indicate “narrow shoulders” where spring and fall months rapidly decline to winter like temperatures.

### Associations between gene expression modules and climate at multiple scales

Co-expression network analysis using WGCNA (44) defined 22 gene co-expression modules (GEMs; mean size: 517 genes per module; range: 54 - 2337 genes; Table S3). 5150 unassigned genes were placed into a dummy module, GEM0, as a useful control for background levels of gene expression divergence. GEMs varied substantially in how strongly their gene expression variation was related to genomic cluster membership (Table S3). For instance, the explanatory value of genomic cluster was rather high for GEM16 (n=665, R^2^=0.20, p<0.0001, Fig. S2a), similar to the unassigned module GEM0 (n=665, R^2^=0.24, p<0.0001), but considerably lower for GEM2 (n=665, R^2^=0.028, p=0.14, Fig. S2b) and GEM6 n=665, (R^2^=0.041, p=0.01). Thus, although background genetic differentiation partially predicts transcriptome-wide expression divergence, expression differences among accessions only weakly reflect co-ancestry for much of the transcriptome.

To assess general relationships between gene expression and climatic variation, we looked for evidence of module expression ~ climate relationships in models including spatial and genomic covariates. Across the 230 climate/module pairs, we identified 23 putative associations (p<0.01, 10x more than expected by chance) with summer precipitation showing the most patterns consistent with climate-associated adaptation in gene expression (Table S4). While these broad associations highlight possible cases of climate adaptation, either due to direct selection on regulation of these gene modules and their downstream phenotypes or due to pleiotropic impacts of selection on other traits, we were particularly interested in parsing the evolutionary scales across which climate adaptation is occurring and assessing the relative consistency or variability of these patterns among clusters of closely related individuals. To do so, we conducted a three-tiered analysis taking into account the geographic scale and consistency of gene expression clines across and within genomic clusters. We label these three approaches as follows: “global”, “consistent among genomic clusters”, and “variable among genomic clusters” (see *Methods).*

#### Global Patterns

By testing for associations between climate variables and cluster mean GEM eigengene values across the 13 genomic clusters spanning the breadth of Eurasian *Arabidopsis thaliana* diversity, we detected likely examples of climate-driven continental-scale adaptation. Using a Storey-Q adjusted p-value correction and an FDR of 0.05 and 0.01, we detected 45 and 28 significant climate-GEM associations, respectively. For example, plants belonging to genomic clusters inhabiting areas with drier summers on average have higher mean expression values for GEM10 (Fig. 3a). Summer precipitation and SSD were the two climate terms that most often explained GEM variation, and the strongest pairwise associations included summer precipitation and precipitation variability with GEM10 (enriched for RNA processing genes), and SSD with both GEM16 (carbohydrate metabolism and cell wall related genes) and GEM18 (glucosinolate metabolism, MYB binding domains) (Tables 1 and S5, Figs. 2b and S3a).

**Figure 3.**
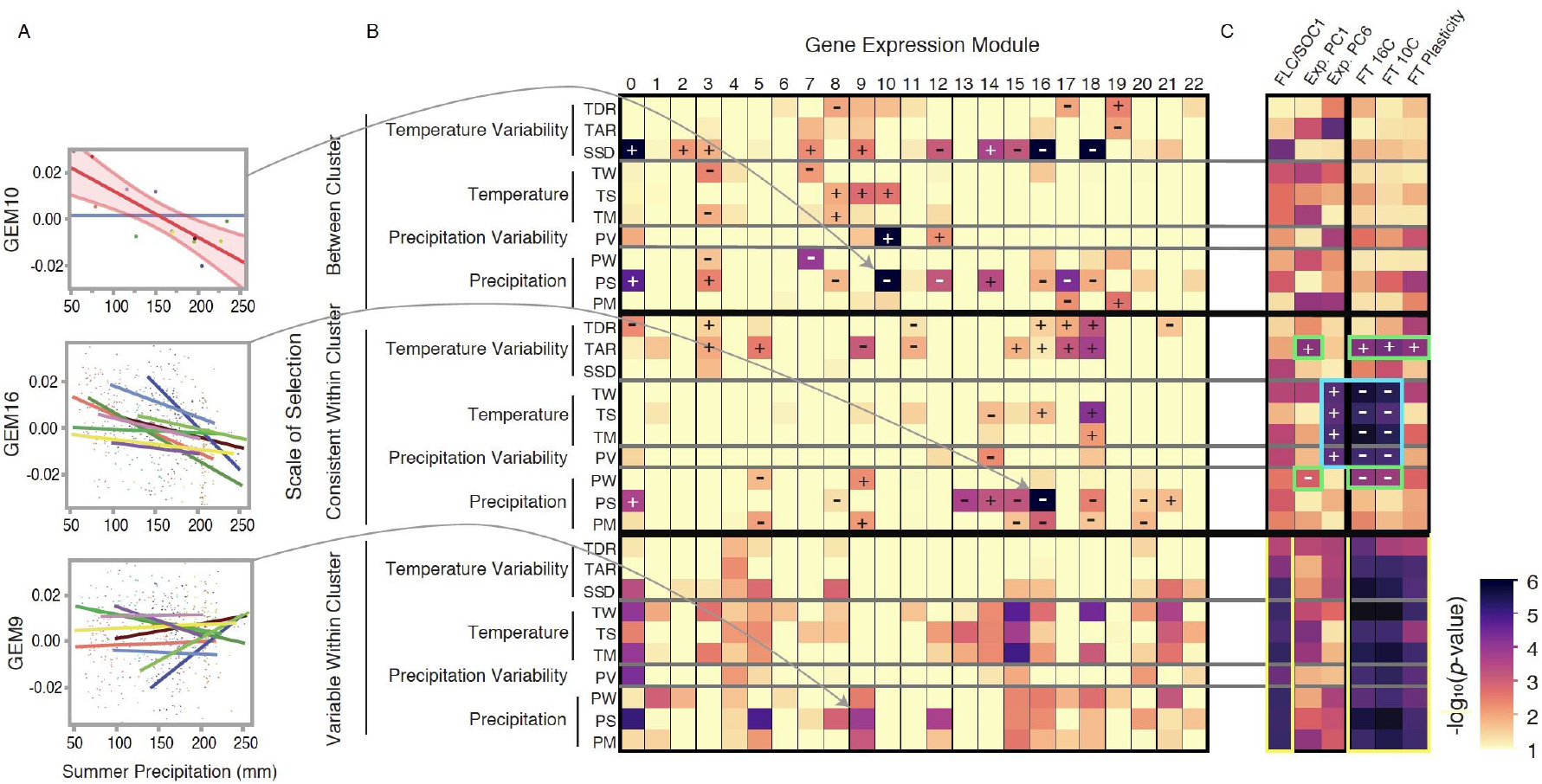
Associations between gene expression modules (GEMs) and climatic variables at three scales. (A) Representative module ~ climate associations across our three scales of selection. Between cluster associations between GEM10 and summer precipitation are identified by the significant negative correlation between genomic cluster mean GEM10 expression level and cluster mean summer precipitation. Associations for all individuals between GEM16 and summer precipitation are consistently negative after taking between cluster expression differences into account, while different genomic clusters of *Arabidopsis* show variable associations between GEM9 expression and summer precipitation. (B) Heatmap showing the log-significance of GEM expression ~ climate associations across three scales of selection. (C) Associations between climate terms and flowering time related sets of genes as well as flowering time phenotypes. Green boxes highlight flowering time ~ climate associations driven by ResPC1 climate associations, blue with ResPC6, and yellow with FLC/SOC1 expression. + : significant positive association, − : significant negative association. Climate term abbreviations defined in the methods section.

**Table 1.** Top climate-associated GEMs (gene co-expression modules) across different scales of selection and enriched categories. Genes: number of genes in the module, Climate: associated climate term and (direction of association), Top MM: Gene with top module membership in the module, Promoter: Most enriched binding site in promoter regions of genes in the module, GO term: most significantly enriched gene ontology term in each module.

| **GEM** | Genes | Climate | Scale | -log(p) | Top MM | Promoter | GO term |
| --- | --- | --- | --- | --- | --- | --- | --- |
| **18** | 93 | SSD (-) | Global | 9.13 | *CB5C* | *MYB92* | glucosinolate biosynthesis |
| **16** | 140 | SSD (-) | Global | 7.7 | *NDL2* | *ARF1* | polysaccharide metabolism |
| **10** | 487 | PS (-) | Global | 6.97 | *ATPEP1* | *NLP4* | RNA processing |
| **13** | 232 | PS (-) | Consistent | 3.71 | *THE1* | NA | cell wall |
| **17** | 107 | TAR (+) | Consistent | 3.21 | *FLN2* | NA | chloroplast |
| **15** | 142 | TM | Variable | 5.22 | *OPR3* | *JAM2* | response to wounding |
| **5** | 674 | PS | Variable | 4.95 | *CNBT1* | *WRKY70* | defense response |

#### Consistent among Cluster Patterns

Next, we tested for significant associations between climate and GEM values using all individual accessions, including genomic cluster as a covariate in the model to account for differences between clusters. A significant association while accounting for clade specific differences suggests that a given climatic factor has consistently driven similar patterns of adaptive divergence in gene expression within groups of related individuals. We detected 33 and 18 significant associations at FDR cutoffs of 0.05 and 0.01, respectively. Summer precipitation and temperature annual range were the climate factors most strongly associated with consistent GEM trends within genomic clusters (Table 1, Figs. 3b and S3a). Negative correlations between GEM13, GEM15 (enriched for jasmonic acid pathway and response to wounding genes), and GEM16 (carbohydrate metabolism, cell wall) eigengene values and summer precipitation are consistently observed within all clusters (Table 1, Fig. 3a), and GEM18 expression is positively associated with summer temperature. These observed relationships between summer climatic conditions and expression of GEM15 and GEM18 (Table 1 and S5), both of which are involved in plant defense, suggest that summer conditions may impact herbivore and pathogen communities, and in turn select for different types and levels of defenses in plants.

#### Variable among Genomic Clusters

Finally, we adapted our models to test for climate x genomic cluster interaction effects on GEM expression levels to assess whether the selection pressure exerted by an environmental factor, aspects of the genomic background, or both have differed among clusters, leading to variability in the magnitude and/or direction of GEM ~ climate associations among genomic clusters. We detected 188 and 81 significant associations at FDR cutoffs of 0.05 and 0.01, respectively. For instance, summer precipitation is positively correlated with mean GEM9 eigengene expression within four genomic clusters, but within three others, these two variables are negatively correlated (Fig. 3a). The most significant climate x cluster interaction terms involved winter temperature, summer precipitation, and mean temperature (Fig. S3a). However, unlike the previous two association categories where adaptive differentiation signals were mostly limited to a few climate variables, we found variable associations to be more evenly spread across climate terms (Figs. 3b and S3b). These results demonstrate that while certain climatic factors drive predictable patterns of adaptive differentiation in gene expression, in many cases locally adaptive signatures are only observable in a subset of genomic backgrounds. GEM4 (enriched for defense genes), GEM5 (response to external biotic stimulus), GEM15 (jasmonic acid), and GEM21 (photosynthesis) show the most evidence of variability in climate-associated differentiation in gene expression patterns (Tables 1 and S5, Figs. 3b and S3b).

#### Lack of overlap across scales and covariate confirmation

Only 8 of 220 GEM ~ climate model combinations showed significant evidence of selection at two of three scales (Table S6), and none showed evidence at all three scales. Thus, it is more the exception than the rule that climate/module associations span scales of relatedness. We did not detect any relationship between the log(p-value) of general module ~ climate associations using latitude, longitude, and genomic PC1 and 2 covariates with our *between cluster* model results (p=0.77). This result reflects how large-scale patterns of climatic and expression differentiation are correlated with spatial and genetic covariation, limiting the capacity of traditional environmental association approaches to identify signals of local adaptation that drive divergence between clades or ecotypes. On the other hand, evidence of selection at the *consistent among cluster* scale is highly correlated with patterns detected using the traditional spatial and genomic PC covariate approach (R^2^=0.51, p<0.001). While there was a significant positive relationship between *variable among cluster* associations and the covariate model (R^2^=0.11, p=0.002), the correspondence was much weaker than for the consistent patterns. This result confirms that associations detected by traditional environmental association analyses will tend to identify adaptive signatures that operate consistently across multiple clinal instances in nature. From a statistical perspective this is reassuring, as this natural replication suggests that the probability of such associations happening by chance or drift are unlikely, but from a biological perspective, it remains unclear what proportion of adaptive differentiation is likely to reflect parallel evolution at this scale of organization.

### GWAS identifies major locus explaining climate-associated gene expression variation

We next took a genome-wide association study (GWAS) approach to identify regions harboring genetic variation that explain eigengene values for the seven climate-associated modules highlighted in Table 1. No significant eQTLs were detected for four GEMs. However, a single ~10kb genomic region (Chr5:22595692-22605170) spanning four genes emerged as an eQTL significantly (FDR<0.001) explaining expression variation for the three defense-related modules: GEM15, GEM16, and GEM18 (Fig. S4). Remarkably, genotype at this interval significantly explains expression variation for 7,338 of the 16,530 genes in our filtered gene set (p<0.01 after accounting for genomic cluster and lat/long effects, enriched in modules 15, 16, and 18, and also 1, 9, and 14), indicating this locus plays a broad role in causing genetic variation in gene expression levels at bolting. The gene set with the 1,000 strongest gene expression associations to genotype at Chr5:22599579 (Table S7) is strongly enriched for glucosinolate-related genes (observed: 15, expected 1.5, p=2.06×10^−6^) and genes involved in blue light responses (observed 18, expected 3.4, p=1.8×10^−4^). Our results are consistent with recent work showing this region harbors an eQTL for 33 flowering time genes, in particular genes annotated as related to flowering time via light-mediated, developmental, or photoperiod control pathways (42).

Although relatively high LD among SNPs across this region prevents pinpointing a causal gene, *AT5G55835* (miR156h) is a candidate for a “master regulator” (first suggested in (45)), as miR156 family microRNAs regulate age-related shifts in glucosinolate and flowering-related expression programs through altering the expression of *SQUAMOSA PROMOTER BINDING PROTEIN-LIKE (SPL)* transcription factors (45). Alternatively, while not particularly well-studied, expression of nearby *AT5G55840,* a pentatricopeptide repeat (PPR) protein, is strongly correlated with the SNP at Chr5:22999579 (R^2^=0.28). Future studies to dissect the molecular basis of this eQTL will be necessary to determine how the highly influential variant(s) in this region achieves such a transcriptome-wide impact at the developmental stage sampled and why this variation helps adapt plants to their local environments.

### Topological Bias of Climate-Associated Gene Expression Patterns

How a gene functions within a gene network is expected to influence how much and what type of genetic variation it harbors (46). Genes annotated as involved in RNA/protein/nucleic acid binding processes are enriched in the top 10% of genes with highest module membership (i.e., similarity to the module eigengene, p<0.0001) within each GEM, indicating these genes likely serve core regulatory functions.

We observed a topological bias in terms of the module membership for genes that showed the strongest patterns of climate-associated differentiation in gene expression, and this bias varied depending on the scale of selection. On average, a gene’s module membership was positively correlated with its maximum *consistent among cluster* log-significance with a climate term (p=0.03) but negatively correlated (p<0.0001) with its *variable among cluster* climatic log-significance. Genes with consistent patterns of climate-associated expression tended to be more highly connected [Module membership of genes with log(consistent selection/variable selection) > 1 = 0.61] than genes with significant climate associations that varied among genomic clusters [Module membership of genes with log(consistent selection/variable selection) < −1 = 0.58), p<0.0001]. Thus, more highly connected genes often contribute to parallel adaptive trends in multiple groups of related accessions, but cluster-specific climate-driven differentiation more often involves the evolution of peripherally connected genes. Des Marais *et al.* (33) recently examined how the plasticity of gene expression to cold and drought stresses varies among *A. thaliana* accessions. They found that the genes with the greatest variability in drought-responsive expression (eGxE) tended to be located in network peripheries (yet, the reverse was true for cold eGxE genes, 26), and here we find a parallel result that genes that have variable associations between gene expression and climate tend to be peripheral, while genes with expression patterns consistently associated with climate tend to be more highly connected.

GEMs 9, 14, 16, and 18 stand out as modules where highly connected genes have more consistent climatic associations than peripheral genes. For instance, the most significant signatures of local adaptation within GEM18 were for highly connected genes negatively associated with summer precipitation (Fig. 4a) and positively associated with temperature annual range. The three GEM18 genes whose individual expression levels are most significantly associated with summer precipitation *(GSTU20, BCAT3,* and *IIL1)* have extremely high membership in this module (Fig. 4b), are involved in glucosinolate biosynthesis, and their expression is higher in plants from drier areas. These findings are consistent with previous work demonstrating that *BCAT3* expression is drought-inducible, that *bcat3* mutants have impaired drought tolerance (47), and that all three genes are regulated by IAA/AUX genes in response to drought stress (48). Drought-inducible gene sets from two independent studies (49, 50) are also significantly over-represented in GEM18 compared to background module GEM0 (ChiSq=10.86, p=0.00098, and ChiSq=12.73, p=0.00036 respectively); 50 of the 93 genes assigned to GEM18 were drought-induced in one or both studies (Table S8).

**Figure 4.**
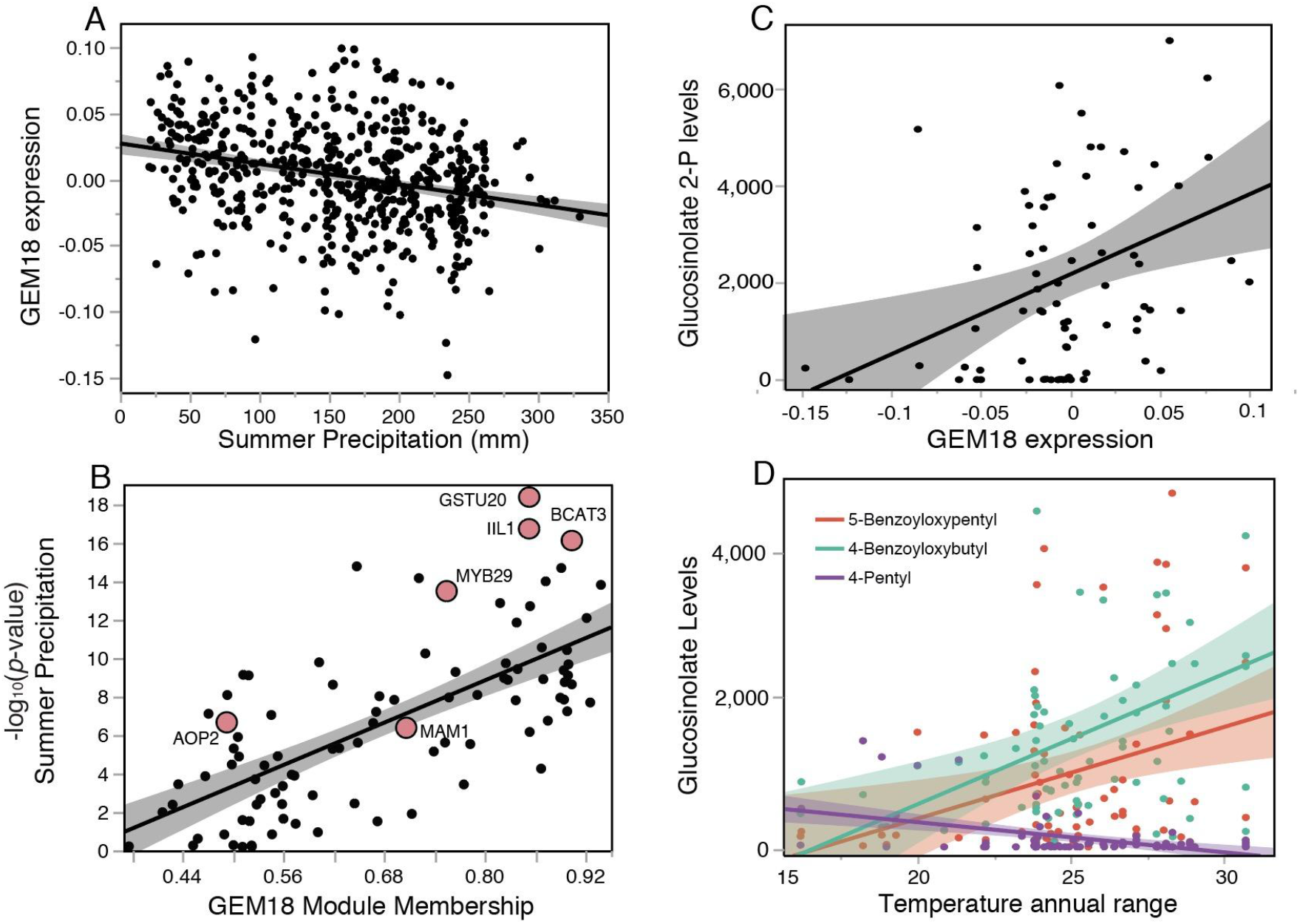
Associations between GEM18 and summer precipitation across 595 accessions. (A) Negative association between summer precipitation and GEM18 eigengene expression level (n=595, R^2^=0.07, p<0.0001). Plants from drier areas tend to express genes in this module more highly. (B) Within GEM18, there is a strong positive association between an individual gene’s module membership value and the log-significance of the association between its expression with summer precipitation (n=93, R^2^=0.47, p<0.0001). Well studied glucosinolate related genes in this module are highlighted and labeled. (C) GEM18 eigengene expression values are strongly associated with plant glucosinolate 2-P levels (n=82, R^2^=0.09, p=0.0049). (D) Associations between temperature annual range and glucosinolate levels in plants are significant and variable, suggesting that different temperatures favor evolution of contrasting batteries of glucosinolates (5-BOP: R^2^=0.10, p=0.0047; 4-BOB: R^2^=0.17, p=0.0001; 4-P:R^2^=0.11, p=0.0027).

### Gene expression variation highlights local adaptation of plant defense traits to climate

Given that several climate-associated GEMs were enriched for genes involved in plant defense, we tested whether these GEMs explained variation in related traits. We first assessed whether GEM eigengene expression could explain variation in trichome density or the presence of colony forming bacteria on plant leaves, as measured in other studies (51). Although the number of accessions for which there is both gene expression and defense phenotype data is relatively low (n=24), noteworthy relationships did emerge between the expression of GEM4 and plant trichome density (R^2^=0.42, p=0.0006, Fig. S5a), and between GEM15 and bacterial titer (R^2^=0.16, p=0.05, Fig. S5b).

The enrichment of GEM18 for glucosinolate pathway genes, its expression differentiation patterns associated with temperature and precipitation, and its enrichment for MYB transcription factor binding sites (p=1.46×10-6) within regulatory regions prompted us to investigate whether variation in glucosinolate levels (52) is explained by GEM18 gene expression levels or these climate factors. Indeed, we detected significant positive and negative associations of GEM18 eigengene expression with one (sinigrin) and three (4-MTB, 4-MSB, and 5-MSP) aliphatic glucosinolates, respectively (Fig. 4c). Notably, GEM18 includes two genes, *ALKENYL HYDROXALKYL PRODUCING 2 (AOP2)* and *METHYLTHIOALKYLMALATE SYNTHASE 1 (MAM1),* that are found in primary glucosinolate QTLs (53) and whose individual expression levels are negatively correlated with summer precipitation, and 19 additional genes with described functions in the glucosinolate pathway are in GEM18 (Table S9). As expected, we detected highly significant positive associations between *AOP2* expression and sinigrin levels (R^2^=0.25) and between *AOP3* expression and 3-hydroxypropyl levels (R^2^=0.33). Likewise, expression of these two genes was negatively correlated (R^2^=0.45), consistent with their roles in shifting flux between alkenyl and hydroxyalkenyl glucosinolates. Six (of 22) glucosinolates showed significant associations with temperature annual range in a minimum BIC selected modules including genomic cluster and longitude (three shown in Fig. 4d). Together, these results strongly suggest that local adaptation to climate has driven differentiation in glucosinolate profiles, in part through modifying expression of members of GEM18.

GEM5, another module that showed intriguing climate-associated gene expression patterns, was also highly enriched for defense-related genes (observed: 92, expected: 18.9; FDR=5.04×10^−30^) and for genes with promoters containing WRKY binding sites (FDR=4×10^−4^). Expression of this module showed the most significant cluster by summer precipitation interaction (Fig. 3b), suggesting that how selection by local climate affects this defensive suite varies substantially across the range of *Arabidopsis.* Consistent with this result, we find a highly significant genomic PC2 x summer precipitation effect on GEM5 expression after taking the main effects of latitude, longitude, and genomic PC1 and PC2 into account (p=0.002). Recent work on Swedish *A. thaliana* accessions described genetic covariation between flowering time and defense-related gene expression (54). Interestingly, we also detected a cluster x GEM5 interaction on days to flower in both 16°C (p=5.0×10^−5^) and 10°C (p=1.7×10^−4^), and these associations are driven by extremely significant positive correlations between GEM5 and flowering for accessions derived from sites above 45°N that are not observed for more southerly accessions (latitude*GEM5, p<0.0001, Fig. S6). This finding highlights the utility of using regional panels to discover patterns of local adaptation but also the caution that trends observed in regional patterns may not reflect range-wide processes.

Recent work has demonstrated that although the positive covariance between flowering time and plant defense and immunity genes is widespread, it can also be decoupled through recombination, suggesting that this predicted tradeoff may emerge more as a consequence of indirect or unrelated selective mechanisms than direct genetic constraints (54, 55). A mutant analysis for 20 *Arabidopsis* transcription factors involved in the aliphatic glucosinolate pathway also inferred a complex relationship between flowering and defense (56), as many mutants affected both glucosinolate levels and flowering time but those effects were not correlated across genes. Together, these results suggested to us that further analysis of this gene expression variation data set in conjunction with climate data could shed new light on the subsets of the transcriptome that contribute to adaptive variation in flowering time.

### Modeling flowering time variation

We made use of the extensive transcriptomic dataset to consider how genetic and environmental variation yield phenotypic variation. Specifically, first taking each data source individually and then evaluating them in combination, we iteratively constructed models to explain variation in flowering time under controlled 16°C and 10°C conditions (FT16 and FT10, respectively) and variation in plasticity of flowering time to these two temperature conditions (FTP) for 597 *Arabidopsis* accessions also represented in the 1001 Genomes genomic and transcriptome data.

#### Insights from Climate

For both temperature and precipitation, annual, summer, and winter mean values individually explain minor (1%-11%) but significant (p<0.01) amounts of variation in flowering time under growth chamber conditions (Table S10a). However, multivariate models including both seasonal and annual terms (3 term models) explain more variance than the sum of the percent variance explained by each term independently (Temperature: 25% of the variation in FT16, 30% in FT10, and 18% in FTP; Precipitation: 14% in FT16, 7% in FT10, and 17% in FTP; Figs. 4a and S6, Table S10a). Significant positive effects of summer and winter temperatures on days to flower, but negative effects of mean annual temperature prompted us to utilize the SSD metric to interpret this result (see *Methods).* A negative effect of SSD on flowering time in models that include mean annual temperature and spatial and genomic covariates reveals that plants derived from areas where fall and spring temperatures are closer to summer than to winter temperatures flowered more rapidly. Similar modeling results with precipitation suggest that the seasonality of temperature and precipitation variation might be a stronger selective force than the mean values of any one term, and that climate seasonality is likely a major player in locally adaptive patterns of phenology. Consistent with previous work (57), we find that a metric that quantifies the timing of seasonal drought explains minor but significant variation in FT16 (4%), FT10 (8%), and FTP (2%). Inclusion of this metric in the multi-term precipitation model increases fit by 5% for FT16, 10% for FT10, and 1% for FTP, suggesting that the timing of seasonal drought may act as a selective agent that shapes flowering time variation, particularly in cool conditions, independent of selection imposed by mean precipitation patterns.

After applying a minimum BIC model building approach to predict flowering time using climate terms and their interactions, we arrived at a set of final models that explain 49% (14 terms), 51% (14 terms), and 34% (12 terms) of the variation among accessions for FT16, FT10, and FTP, respectively (Fig. 4a and 4b, Table S10b). Previous common garden studies in *Arabidopsis* have differed in whether or not they detected flowering time clines with latitude or climate gradients (58–60). We predicted this discordance reflects regional differences in adaptive differentiation, and indeed, applying the three-tiered framework used for the GEMs above, we find that FT16, FT10, and FTP all show many clinal signatures of local adaptation to climate that vary among genomic clusters (Fig. 3c). For example, for accessions belonging to genomic cluster 3, lines with source sites that experience warmer winters flower earlier, but in contrast, accessions belonging to genomic cluster 6 show the opposite pattern, and cluster 9 accessions show no flowering time trend with winter temperature (Fig. S7). Overall, our findings reinforce the conclusion that many signatures of natural selection may operate over relatively fine environmental grains and that local adaptation can often generate distinct relationships between phenotype and climate in different regions.

#### Insights from Genomic Variation

To model the influence of genetic variation on flowering, we first examined whether genomic ancestry and flowering time covaried and found that genomic cluster membership explained 32%, 34%, and 25% percent of the variation in FT16, FT10, and FTP, respectively (Figs. 5a and S8). Next, we built models using sets of flowering-time associated SNPs previously identified through GWAS or other approaches (61). Models based on these flowering-time SNPs explained between 40% and 66% of the variation in FT16, FT10, and FTP (Fig. 5a). However, because the numbers of SNP terms in these models present a significant overfitting problem, we applied a model selection approach to identify reduced sets of SNPs that explain flowering time variation after including genomic background terms (Fig. 5a). The resulting models reduced the number of included SNP terms from 23, 25, and 37 SNPs to 7, 6, and 3 SNPs for the FT16, FT10, and FTP models, respectively, with only minor reductions in the variance explained by the model (≤ 5.1%; Figs. 5a and S8). These refined models including a limited set of SNPs and genomic cluster information explained 55%, 57%, and 36% of the variation in FT16, FT10, and FTP, respectively (Figs. 5 and Table S11).

**Figure 5.**
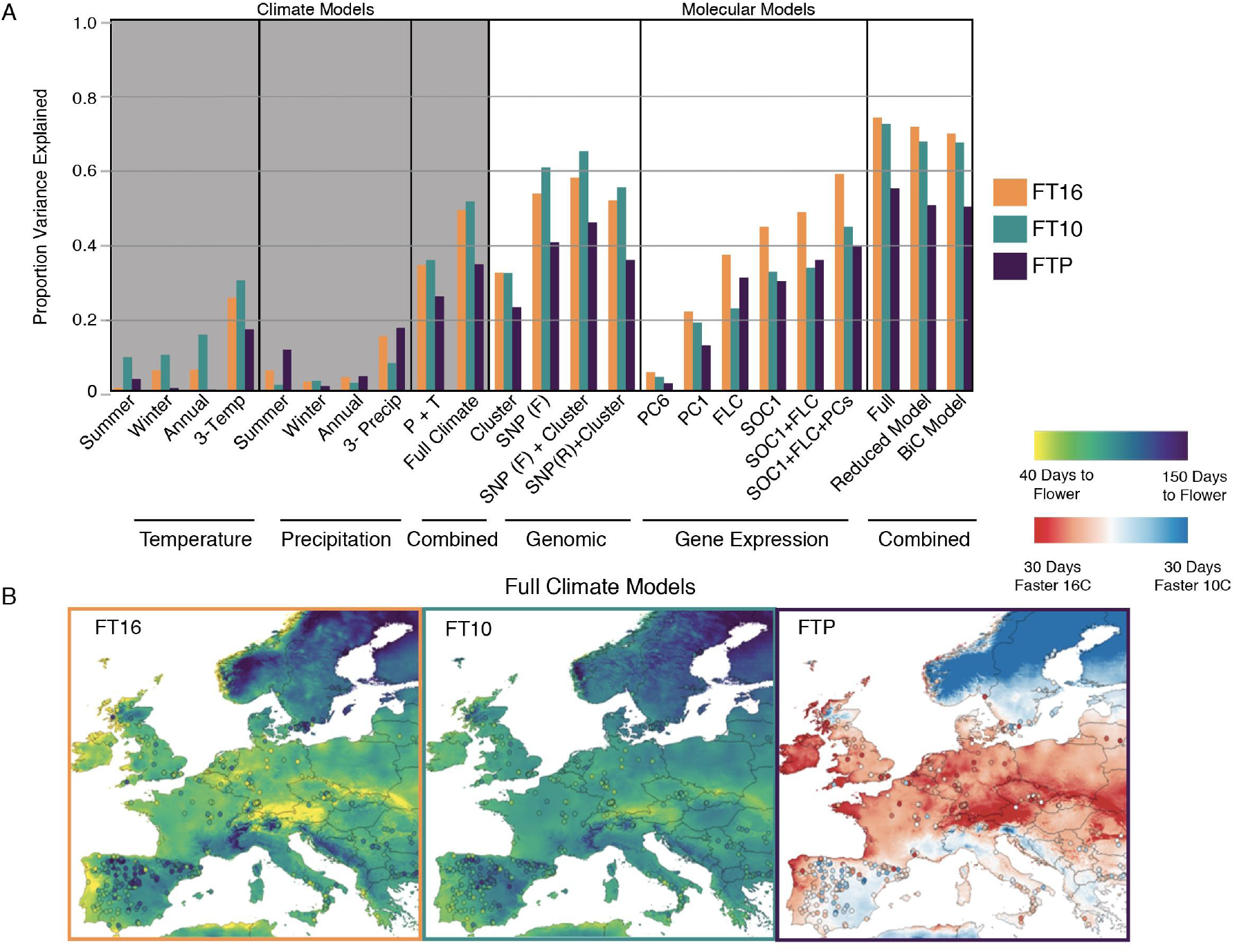
Comparison of models constructed to explain variation in flowering time. (A) Bar plots of the proportion variance explained by models constructed using climatic or molecular terms. SNP (F): Full set of SNPs, SNP(R): Reduced set of minimum BIC selected SNPs, (P + T): Models including both precipitation and temperature terms. (B) Maps presenting the full climate model applied to raster climate terms across the study range. Points colored to show individual *Arabidopsis* accession phenotype values.

#### Insights from Transcriptomic Variation

We next developed models to predict flowering time from gene expression data. Individual gene expression levels were correlated with flowering time for a large fraction of the transcriptome (FT16: 3921 correlated transcripts, FT10: 3811 correlated transcripts; FDR-adjusted p-value < 0.01). Since many genes are co-expressed, we further parsed these gene sets to identify strong predictors. MADS-box transcription factors *SUPPRESSOR OF OVEREXPRESSION OF CONSTANS 1 (SOC1)* and *FLOWERING LOCUS C (FLC)* were the genes whose transcript levels were most highly correlated with flowering time, explaining 44% and 36% of the variation in FT16, respectively (Fig. 5a). A model including both terms only has marginally increased fit (48% for FT16) because these two predictors were highly correlated, consistent with *FLC’s* function as both a direct (62) and an indirect (63) negative regulator of *SOC1.* A substantial subset of the transcriptome covaried in expression with *SOC1* and *FLC* expression; >10% of the expression variance for 1,184 genes was explained by the levels of these two transcription factors (p<0.0001, Table S12). Of these, 650 genes had transcript levels positively associated with *SOC1* expression (and negatively associated with *FLC* expression), and 534 genes showed the opposite pattern.

#### Genes positively associated with SOC1 expression

GEM2 and GEM7 were heavily enriched for transcripts positively associated with *SOC1* levels. Although no GO terms are enriched among these GEM7 transcripts, the *SOC1-*associated transcripts in GEM2 are enriched for response to temperature and circadian rhythm GO terms. The binding site motifs most highly enriched in the promoters of these genes were those bound by circadian clock related transcription factors *(LHY, RVE6, RVE8* and *RVE1)* and the CArG box motif (CC(A/T)6 GG) bound by MADS-box transcription factors like *FLC* and *SOC1.* The expression of *PRR5,* a core clock gene (64), is highly associated with *SOC1* expression (p=7.36^−14^, R2=0.15), and it also has the fifth highest module membership (0.932) for GEM2. Consistent with enrichment for their binding motifs in GEM2, *PRR5* directly represses expression of *LHY* and *RVE* genes across a broad diversity of angiosperms (65). In addition, *PRR5* directly represses three *CYCLING DOF FACTOR* transcription factors known to repress flowering by inhibiting transcription of *CONSTANS,* which directly and indirectly activates *SOC1* transcription (66–70). Thus, natural variation in *SOC1* expression may derive both from variation in the expression of factors that promote its expression as well as factors that repress its expression (i.e., *FLC).*

GEM2 was also highly enriched for genes upregulated as *A. thaliana* plants age (54)(146/385 gene up-regulated at 28 d vs. 14 d post-germination in Bur-0; FDR <1.0×10^−5^). In addition, the first principal component of variation in an analysis of 612 genes differentially expressed between young and old plants in this same experiment (age PC1) was highly correlated (r=0.93) with GEM2. A structural equation model with *SOC1* impacting both days to flowering and the expression of age PC1, and in which days to flower also impacts age PC1 through an “age at bolting” latent factor (X^2^ from independent =451.62, SRMR=0.007, CFI=0.99, Fig. S9) best fits the data. These results suggest that *SOC1* expression induces rapid flowering as well as plant aging through partially distinct mechanisms. Recent work has demonstrated that *SOC1/FUL* double mutants not only have extremely delayed flowering, but actually transition to a perennial growthform (woody stems, bushy growth form, etc.), indicating that *SOC1* functions to regulate developmental processes in addition to the floral transition (71).

#### Genes negatively associated with SOC1 show latitude-dependent associations with flowering time

Members of two co-expression modules with significant climate associations, GEM5 (defense-related) and GEM16 (carbohydrate- and cell wall-related), were enriched in the set of transcripts whose expression is negatively associated with *SOC1* expression. We detected a consistent negative association between GEM16 and *SOC1* expression across genomic clusters (R^2^=0.2, p<0.0001), but as described above, GEM5’s association with *SOC1* and *FLC expression* are highly cluster dependent (GEM5 ~ *FLC\**cluster: p=0.00007). These results suggest that while the expression of genes in GEM5 is associated with the expression of these core flowering time genes, other factors likely exist that act in a latitude-dependent fashion to modify this relationship between GEM5-related defense traits and flowering time.

#### Gene expression patterns associated with flowering time independent of SOC1/FLC

We next identified a set of 260 genes whose expression levels are significantly associated with FT16, after controlling for *SOC1/FLC* expression levels (FDR<0.01). To reduce the complexity of this additional gene set, we conducted a principal component analysis and screened the resulting PCs. Expression PC1 and PC6 were the axes of gene expression variation most highly correlated with flowering time across the diversity panel, explaining 16% and 4% of the variation in FT16 and FT10, respectively (Fig. 5a). Adding these two PC values as effects to a flowering time variation model with *SOC1/FLC* expression increased the explanatory power from R^2^=0.48 to R^2^= 0.58 (Fig. 4a). The set of genes with strong loadings on expression PC6 were not enriched for any GO terms or GEMs. In contrast, expression PC1 values are highly correlated with GEM5 eigenvalues (defense related; R^2^=0.60). Notably, *WRKY70,* a key mediator of cross-talk between the JA- and SA-signaling defense pathways (72), was the binding motif most enriched in GEM5 promoters and the third highest contributor to expression PC1 loading, and genes with strongly negative (<-0.3) loadings on PC1 (Table S13) were enriched in the phenylpropanoid pathway (p=1.5×10^−5^). Although either overexpression or antisense suppression of *WRKY70* has been found to alter flowering time (72), how this functions in the flowering time regulatory network has been examined. These findings indicate that *WRKY70* may function as a key transcriptional integrator of upstream signals like *FLC* and *SOC1,* and suggest that adaptive modulation of its expression may facilitate co-evolution of plant defense and flowering traits.

#### Separable components of flowering-associated gene expression relate to climate at different scales

We next assessed how each of these three major components of flowering-associated gene expression variation--*FLC/SOC*-related expression, expression PC1, and expression PC6--are distributed with respect to climate and genomic clusters. *FLC/SOC1* expression showed significant associations with all ten climatic factors, but these associations mainly varied among genomic clusters (Fig. 3c). In contrast, we found significant associations between winter temperature or precipitation with expression PC1 values that were consistent across genomic clusters, suggesting that tradeoffs between defense and flowering time may be related to adaptation to winter conditions (Fig. 3c). The distinct relationships of different flowering-associated gene expression axes with climate parameters is well illustrated in the case of mean annual temperature (Fig. S10). Expression PC6 was consistently positively associated with mean annual temperature across clusters, while the nature of the relationship between this climate term and *SOC1/FLC* was highly cluster-dependent. Cumulatively, the patterns of expression differentiation correspond to all 16 of the significant climatic correlations found for FT16 and FT10 (highlighted boxes in Fig. 3c), indicating that both range-wide and cluster-specific patterns of flowering time adaptation arise from these three major axes of expression variation. Thus, different and distinguishable underlying molecular phenomena have been harnessed to promote flowering time adaptation at different geographic scales in response to local climatic pressures.

#### Insights from Integrative Flowering Time Models

The models we have developed thus far examine how variation in different types of factors--climate parameters, genomic ancestry, trait-associated SNP genotypes, and gene expression--individually explain natural variation in flowering time and its plasticity to temperature. However, several of these factors co-vary. For instance, variation in gene expression patterns associated with flowering time and climate adaptation should derive at least in part from allele frequency clines at known flowering-associated SNPs. Therefore, to develop an integrative understanding of how these individual components of variation jointly shape natural variation in flowering time, we applied a sequential model building approach that explicitly tested the extent to which gene expression data provides explanatory value above and beyond the information content of previously identified flowering-associated SNPs. When major gene expression and genotypic effects are included in the model with model selection based on 5-fold cross-validation, the portion of natural variation in FT16 explained rises to 69% (d.f. = 21), compared to 55% for either SNP or 58% for gene expression data alone (Table 2, Table S14, Figs. 5a and S8). Likewise, a joint model including genotype and gene expression explains 69% of the variation in FT10, compared to 57% for the genotype model and 44% for the gene expression model alone (Table 2, S13). We note that inclusion of gene expression data enhances the predictive value of both models even though gene expression measurements were obtained from plants grown at 22°C. To further validate our finding that gene expression variation accounts for flowering time variation distinct from genotypic information, we also tested whether *SOC1* expression, *FLC* expression, or either expression PC1 or expression PC6 (residual gene PC axes after accounting for *SOC1* and *FLC*) explained additional variance after accounting for genotypic variation in larger sets of all 23 previously identified flowering-associated SNPs. After factoring out all the variation explained by either SNP panel, we find that *SOC1* (13.5%, p<0.0001), *FLC* (8.3%, p<0.0001), and PC6 (1.5%, p=0.049), but not PC1 (p=0.14) still explain significant portions of residual FT16 variance.

**Table 2.** Minimum BIC selected molecular models of flowering time variation.

| Molecular Model |  |  | 16C Days to Flower (n=412, df=21) |  |  |  | 10C Days to Flower (n=430, df=20) |  |  |  | FT Plasticity (n=544, df=18) |  |  |  |
| --- | --- | --- | --- | --- | --- | --- | --- | --- | --- | --- | --- | --- | --- | --- |
| Class | Scale | Source | Effect | SS | F-Ratio | LogWorth | Effect | SS | F-Ratio | LogWorth | Effect | SS | F-Ratio | LogWorth |
| GE | Ind. Genes | *SOC1* Expression | -5.23 | 27594 | 140.25 | 28.50 | -3.05 | 16313 | 155.00 | 31.20 | -2.1 | 3918 | 43.64 | 10.01 |
|  |  | *FLC* Expression | 0.99 | 974 | 4.95 | 1.58 |  |  |  |  |  |  |  |  |
|  | Exp PCs | PC1 | 0.94 | 9234 | 46.93 | 10.74 | 0.65 | 5434 | 51.69 | 11.72 | 0.47 | 2000 | 22.27 | 5.52 |
|  |  | PC6 | -1.19 | 3149 | 16 | 4.15 | -0.89 | 1839 | 17.49 | 4.48 | -0.43 | 358 | 3.98 | 1.33 |
| Genetic Ind. SNPs |  | 5_18590501 (*DOG1*) | 3.72 | 4521 | 11.49 | 4.89 |  |  |  |  |  |  |  |  |
|  |  | 5_18590327 (*DOG1_2*) |  |  |  |  | 1.63 | 4315 | 20.52 | 8.63 |  |  |  |  |
|  |  | 5_22979827 (*VIN3*) | 7.65 | 1518 | 7.26 | 2.13 | 6.25 | 3288 | 15.64 | 6.62 |  |  |  |  |
|  |  | 2_8137542 (*PHYB*) | 3.65 | 1375 | 6.57 | 1.97 |  |  |  |  |  |  |  |  |
|  |  | 5_26805231 (*TOE3*) | 3.9 | 1773 | 4.5 | 1.94 |  |  |  |  |  |  |  |  |
|  |  | 2_19200447 (*NF-YB12*) | 3.07 | 1539 | 3.91 | 1.69 |  |  |  |  |  |  |  |  |
|  |  | 4_10999188 (*TSF*) |  |  |  |  | -3.75 | 5067 | 24.09 | 10.08 |  |  |  |  |
|  |  | 1_24339560 (*FT*) |  |  |  |  | 1.08 | 1108 | 5.27 | 2.27 |  |  |  |  |
|  |  | 2_16387379 (*SNZ*) |  |  |  |  | -0.31 | 734 | 3.49 | 1.51 |  |  |  |  |
|  |  | 4_17882277 (*BHLH131*) |  |  |  |  |  |  |  |  | -2.88 | 3289 | 18.35 | 7.74 |
|  |  | 1_23453207 | 6.09 | 7115 | 18.08 | 7.63 |  |  |  |  | 3.18 | 2589 | 14.97 | 6.93 |
|  |  | 1_7885760 |  |  |  |  | 2.92 | 3508 | 16.69 | 7.06 |  |  |  |  |
|  |  | 4_17282047 |  |  |  |  | 3.04 | 1967 | 9.36 | 4.00 |  |  |  |  |
| Cluster |  | Genome Cluster (13) |  | 9852 | 4.17 | 5.59 |  | 6629 | 5.25 | 7.74 |  | 4022 | 3.74 | 4.76 |
|  |  | Model |  | 177874 | 40.49 | **R2=0.690** |  | 93112 | 41.42 | **R2=0.667** |  | 46415 | 28.72 | **R2=0.494** |
|  |  |  | **R2Adj=0.676** |  |  |  | **R2Adj=0.653** |  |  |  | **R2Adj=0.477** |  |  |  |
|  |  |  | **R25-Fold=0.645** |  |  |  | **R25-Fold=0.635** |  |  |  | **R25-Fold=0.457** |  |  |  |

Because gene expression under constant 22°C conditions explained ample variation in flowering and its plasticity under other constant temperatures, we next assessed whether these data can also reasonably inform prediction of flowering time variation for plants raised in more ecologically realistic conditions. For flowering time data collected for Spain and Sweden spring and summer field conditions in both 2008 and 2009, models including gene expression data explained significantly more variation (37% - 73%, mean: 61%) than genotype only models (23% - 59%, mean: 44%, Table S15). In all cases, the FT10 integrative model predicted variation in flowering time slightly better than the integrative FT16 model across the diverse climate-matched experimental conditions (marginal increase from 0.2% to 6.5%). One possible explanation for this observation is that in all four conditions, the daily minimum temperatures dropped to 12°C (or lower) in the evenings, and while all reached above 16°C during the days, the colder 10°C chamber may have better emulated the relevant temperature cues for floral induction.

#### Relationships between terms

To identify related flowering time associated terms, we performed a clustering analysis on the terms included within these models (Fig. S11). This analysis identified 11 clusters of between 2 and 7 terms that covaried, likely indicating shared molecular factors or selective pressures underlying their variation. For instance, our major eQTL SNP along with three previously identified flowering SNPs (61) in this region show highly concordant patterns of variation. Predictably, we also observed high covariance between genotypes at SNPs in *FRIGIDA (FRI)* and *FLC* and *SOC1* and *FLC* expression levels. Genotype at *DELAY OF GERMINATION 1 (DOG1),* which has known impacts on germination and flowering, also grouped with this cluster. The high covariance observed between *DOG1* genotype and these additional core flowering time genes likely reflects a history of correlational selection acting on germination and flowering variation to shape the range-wide distribution of life-history syndromes (73), although undescribed direct impacts of *DOG1* on *FLC/SOC1* expression cannot be excluded.

Next, to account for the hierarchical relationships between climate, genotype, gene expression, and phenotype, we constructed a model using a path analysis approach (Fig. 6). This framework allowed us to visualize the co-variances among spatial, environmental, genotypic, and gene expression flowering time factors while also determining the strength of connections after accounting for other related terms. The resulting model recaptures the finding that genomic PC1 is highly correlated with temperature SSD, providing further evidence that the dynamics of temperature change across the season may be a major factor that structures genetic variation of *Arabidopsis* across Eurasia. Within this framework, we find the causal sequence that explains the largest amount of variance in flowering time leads from latitude through mean temperature then through genotype at *FRI* (SNP 4_259252) affecting *FLC* expression and in turn *SOC1* expression (Fig. 6: red connections), consistent with the well-known impacts of *FRI* alleles on natural variation in *FLC* expression and flowering (59). Additionally, we find a relatively minor, but significant link between annual temperature, genotype at the Polycomb group protein FERTILIZATION INDEPENDENT ENDOSPERM (FIE; SNP_3_724007), and expression PC6 with flowering time, suggesting that flowering time adaptation to temperature might occur through downstream consequences of genotype at this epigenetic regulator (Fig. 6: Blue connections). The association of *DOG1* genotype with both *SOC1* expression and residual PC1 may reflect its complex ABA- and miR156-dependent association with flowering time (74).

**Figure 6.**
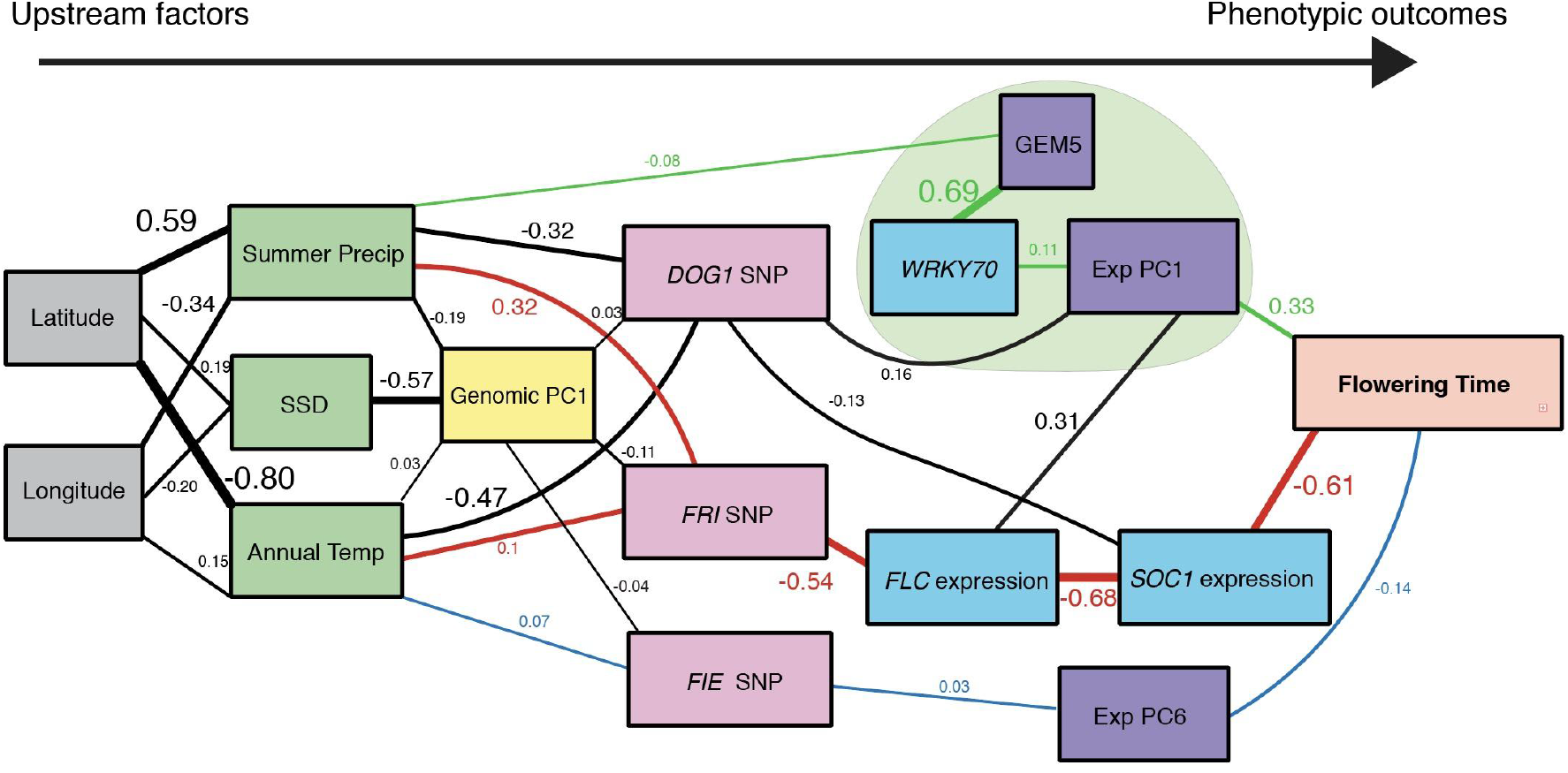
Path analysis showing connections between spatial, climate, genetic, gene expression, and flowering time variation (FT16). All lines represent regression relationships with the explanatory variable left of the response variable, with the exception of the three connected gene expression terms above the green background which are modeled as covariance relationships due to the uncertain arrow of causality. Box colors reflect different classes of data, and numbers above lines and line weights reflect the regression estimate for that path. Green paths follow the link from summer precipitation to gene expression terms associated with defense and explaining residual flowering time variation. Red paths represent the connection between temperature and precipitation with genotype at the *FRI* locus, and in turn *FLC* and *SOC1* expression, which itself explains the largest portion of flowering time variation. The blue lines show the links between temperature, *FIE* genotype, and residual PC6 with flowering time.

Lastly, a substantial amount of flowering time variance is explained by summer precipitation shaping adaptive differentiation in the expression of GEM5 and *WRKY70,* which in turn appears to mediate the regulation of genes involved in a tradeoff between flowering time and plant defense (Fig. 6: green connections). As an integrator of various upstream signals related to pathogen, herbivore, and drought signals, *WRKY70* plays a key role at the intersection of the JA, SA, and brassinosteroid pathways (72, 75). While its role in plant defense has been studied extensively (72, 76), recent work has provided evidence that it also has a more widespread role in regulating plant growth (75). In addition, its expression is regulated by histone variant H2A.Z, an epigenetic modifier that also plays a large role in FLC-mediated epigenetic floral repression (77) and in regulation of *FLOWERING LOCUS T* by ambient temperature (78). These results suggest that selection by local precipitation regimes may have shaped the evolution of *WRKY70* regulation, such that expression of this signal integrator may have been adaptively fine-tuned to allocate resources between abiotic/biotic stress tolerance and plant growth/reproduction.

## Conclusion

By individually and jointly analyzing large-scale datasets describing genetic, transcriptomic and flowering time variation, we reveal substantial new insight into how local adaptation to climate has proceeded historically at multiple evolutionary scales and levels of organization in *Arabidopsis thaliana.* Our results demonstrate that climatic variation has likely driven adaptive differentiation of several gene co-expression modules. Although some of these signatures are consistent among different relatedness groups on a continental scale, many other patterns are region-specific and not adequately captured by solely applying standard statistical approaches for detection of trait-environment associations or with locally intensive sampling (12). Notably, we find that genes whose expression patterns display associations with climate parameters consistent across genomic clusters are more highly connected within co-expression modules than genes whose expression pattern associations with climate vary from region to region. One possible explanation for this pattern may be that variation in genes with greater connectivity often involves functional tradeoffs of adaptation to co-varying environmental pressures. In contrast, variants in peripheral genes may be less constrained by pleiotropy or more often have substitutable phenotypic effects, thus providing a greater diversity of evolutionary paths to achieve adaptation to regional conditions.

In addition to studying how climate-mediated selection has shaped patterns of gene expression differentiation as a molecular phenotype, we also investigate how gene expression can serve as an informative intermediate phenotype for connecting climatic variation with adaptive evolution of observable organismal phenotypes. For instance, our analyses call attention to plant defense evolution as a fundamental contributor to local adaptation in this species, as climate-associated patterns of gene co-expression highlighted adaptive differentiation in glucosinolate composition potentially due to their joint impacts on drought tolerance and herbivory. The lasting imprint that historical climatic conditions have left on defense-related gene expression networks further underscores the necessity of research that considers how the impact of climate change on plant interactions with pathogens and predators will vary by genotype and region (79). Our approach also allowed us to dissect the distinct climatic factors that shape different gene expression-mediated components of variation in a key life-history trait, flowering time. Consistent with past work (41), we found that the *FLC/SOC1* axis of expression variation explains the largest component of flowering time variation. We also revealed a latitude-dependent relationship between plant defense module expression and flowering time, and that *WRKY70* likely serves as a central integrator in this relationship. Future studies addressing *WRKY70’s* role in mediating relationships between growth, reproductive timing, and defense should shed light on how diverse combinations of selective pressures shape co-variation of these highly multivariate phenotypes across complex landscapes.

Finally, through constructing linear and hierarchical models of flowering time variation that incorporated climatic, genomic, and transcriptomic data, we revealed the organization by which distinct components of expression variation corresponded to range-wide and region-specific associations between flowering time and climate. In addition, we find that including gene expression data significantly increases the amount of flowering time variation that can be explained compared to a model solely including genetic variant information, potentially because gene expression captures variation resulting from epistatic interactions (56) or epi-allelic variation (80). Moreover, even though these models were informed by gene expression data collected in a single environment different from the conditions under which phenotype data were collected, incorporating this transcriptomic information still improved models of trait variation across multiple ecologically relevant conditions. Thus, our results suggest that even when joint sampling of different data types is not ideal due to the factorial cost of sampling transcriptomes from many genotypes across multiple environments, single environment datasets can still be of great interpretive or predictive value and yield ample information about the molecular and environmental drivers of phenotypic variation.

## Materials and Methods

### RNA-seq data

The normalized RNA-seq read count table produced by the *Arabidopsis* 1001 Epigenomes Project (16) was downloaded from the NCBI Gene Expression Omnibus (GSE80744). This data was generated for each accession from pooled samples collected directly prior to bolting from ten individual rosette plants grown at a constant 22°C in 16h light : 8h dark cycles. We analyzed data for the subset of 665 accessions (of 728 total) for which we could also obtain corresponding SNP data, flowering time data from at least one experiment, and latitude/longitude of origin information.

#### Filtering and WGCNA

We used edgeR (81) to filter out lowly expressed genes and to normalize the data. Only genes with >5 counts per million (cpm) for at least 10% of the individuals were used in gene module construction. Read counts were transformed in limma using the “voom” function (82). Gene expression modules (GEMs) were constructed with WGCNA (44) using a “soft power” of 5 and a minimum of 60 genes per module.

### Genomic clustering and SNP extraction

Working with the unpruned SNP matrix from the 1,135 Genomes project (43), we used SNPRelate (83) to assess relationships between accessions and define genomic similarity clusters. The first seven principal component axes (PCs) were extracted (Table S1). Because we lacked an *a priori* expectation for the number of clusters, we processed the genomic PCs by Ward hierarchical clustering using JMP Pro 14 (84) and then used the cubic clustering criterion metric, which classified individuals into 13 clusters and one unclustered accession that was excluded from subsequent analysis. Using vcftools, we exported SNPs previously associated with flowering time variation (42), climate variation (85), and drought (86) into a binary allele table by individual, retaining SNPs with <40% missing data across accessions.

### Flowering time and defense phenotypes

Previously, flowering time was measured under 16°C and 10°C (43) constant temperature conditions (16h light : 8h dark cycles in both conditions) for the accessions for which we have RNA-seq data (16). Importantly, these conditions differ from the 22°C at which plants were grown for gene expression analysis. Lack of paired data types from the same individuals will weaken some expression ~ phenotype relationships, reducing our power to detect the underlying expression signatures of trait variation, but should not lead to spurious false positive associations. Additionally, we extracted flowering time data from a subset of accessions that were grown in chamber conditions designed to mimic field conditions in Sweden and Spain (87). Previously obtained trichome density, bacterial titer, and glucosinolate trait data were acquired for a subset of accessions (51).

### Climate Analysis

#### Climate associations with genomic clusters

We downloaded all 19 WorldClim bioclim layers (bioclim 2.0) (88) at 30-second resolution and used the QGIS point sampling tool to extract climate data for all accession source site coordinates. We determined the climatic factors most highly associated with genomic clusters by nominal logistic regression. We compared a model only considering spatial distance (lat./long.) with one including both spatial distance and each of the ten possible climate factors. To develop an integrative climatic model for genomic differentiation, we iteratively added the most highly significant climate terms until model BIC began to increase.

#### Seasonal Sine Deviance

Initial exploratory analyses revealed that models incorporating multiple seasonal temperature terms explain more variation in genomic PCs, clusters, flowering time, and gene expression modules. In particular, the direction and extent to which spring and fall mean temperatures deviated from the average of the winter and summer mean temperatures explained most of the variation captured by including each of these terms in the model. Seasonal temperature variation is often formalized as a sine curve where the y-value (temperature) at the time directly intermediate the peak and trough of the curve is the mean of the peak and trough y-values. However, in natural environments, temperatures during the intermediate months often deviate to be more similar to either the summer months or the winter months. We termed this offset from a pure sine curve the *seasonal sine deviance* (SSD), and calculated it as ((T_*spring*_ + T_*fall*_) - (*T_summer_* + T_*winter*_)) / (T_*summer*_ + T_*winter*_).

### Climate associations of gene co-expression modules

For our initial examination of of GEM expression ~ climate relationships, we calculated individual eigengene expression values for each of the 23 GEMs and then fit separate generalized linear models in JMP 14 with these as response variables; each of the 10 climate terms as an explanatory variable; and latitude, longitude, and the first four genomic PCs as covariates (230 tests total). The eigengene expression level captures the first principal component axis of expression variation of all genes assigned to a module. We ran each test with three variants: a traditional generalized linear model, one using the Huber robust method, and a variant using a Cauchy rather than normally distributed error term. To de-emphasize highly significant (low p-value) outliers, we used the geometric mean of the negative log transformed p-values produced by these three approaches as our metric for assessing the significance of association between climate variables and gene expression.

Next, to identify the scale and consistency of gene co-expression module (GEM) ~ climate associations, we tested for significant relationships between module expression and climatic terms at three evolutionary levels. First, to assess signatures of widespread adaptive differentiation across our 13 genomic clusters, we calculated the cluster means for the 10 climate values and for the module eigengene expression levels for the 23 GEMs. We then applied the three modeling approaches (Normal, Cauchy, Huber) as described above to quantify the strength of evidence of association between the climatic conditions that individuals in a genomic cluster tend to be found and the average eigengene expression values.

Second, to look for consistent patterns of local adaptation within, rather than between, genomic clusters, we performed a response screening approach using a subset of 597 individuals filtered to foster joint analyses with flowering time (see below). Specifically, we tested for associations between GEMs and climatic factors while also accounting for differences in expression due to differences between clusters. We fit traditional, Huber, and Cauchy linear models incorporating genomic cluster membership and each of the ten possible climate terms for each of the 23 expression modules using the method described above to evaluate significance.

Finally, to identify within-cluster associations between GEMs and climatic factors that varied in strength or direction among different genomic clusters, we modified the previous traditional, Huber, and Cauchy models to include the interaction between genomic cluster (nominal) and climate term and evaluated the results by calculating the geometric mean of the log-transformed p-values associated with the interaction term. For each of the three scales of association, p-values were loaded into R as a vector and processed with the “qvalue” package to account for multiple testing.

To compare evidence of selection between the three scales measured as well as the traditional co-variate approach, we looked for correlations in the −log(p-values) between types for each of the 230 tests. To confirm the robustness of these results at each of the three scales of associations, we re-ran the same models and included latitude and longitude or included genomic PC1 and PC2 as additional covariates for a subset of four climate terms.

#### Specific genes most closely associated with climate patterns

To determine the individual genes within GEMs that were most closely associated with climate terms of interest and to determine whether these genes show particular patterns of connectivity and mean expression values, we looked for associations between the expression of individual genes and climate terms, considering models that included a combination of genomic cluster, climate parameter, and their interaction terms. We used −log(p-value) as a metric to assess the significance of these associations.

Next, to test for associations between a gene’s module membership value (relatedness of a gene to the eigengene value of that GEM) and the significance of the relationship between its expression and a climate term, we fit a model considering the effect of gene module membership on the −log(p-value) for that gene’s most significant consistent association (climate effect). We then performed the same analysis using the −log(p-value) for its most significant variable association (climate-by-cluster interaction effect). We then calculated the log significance ratio of consistent/variable climate associations. A ratio > 1 indicates the association of a gene’s expression with climate is more consistent across clusters; a ratio < −1 indicates a gene’s association with climate is more variable across clusters. We then performed a t-test to compare mean module membership between these two groups both for all genes and for each module.

#### Promoter motif and GO enrichment analysis

For each gene group tested, we extracted transcript sequences from TAIR database (version Araport11) and used the PlantRegMap Database (89) to perform enrichment analysis of promoter regions 2000 bp upstream of genes. Gene Ontology (GO) enrichment analysis was performed using PANTHER14.1 (90).

### eQTL mapping for climate associated GEMs

We used the easyGWAS web service (91) to conduct eQTL mapping on module eigenvalue levels for seven GEMs with significant climate associations using the first five PCs of genetic variation as covariates and excluding SNPs with MAF < 0.05. SNP information from genomic regions with significant associations was extracted and further analyzed in JMP. To identify GEM members whose expression levels are highly associated with SNPs in the GWAS hit region, we used JMP 14 response screening by including the first five genomic PCs, longitude, and latitude as covariates.

### Associations between genomic cluster, gene expression, climate and flowering time

#### Climate and flowering time

We reduced our dataset from 665 accessions to 597 accessions from Europe or mainland Asia to exclude recently introduced accessions and to leave a maximum of two accessions derived from any one sampling location. After performing standard linear regressions to calculate correlation coefficients between climate factors, we applied a minimum BIC model selection approach with 5-fold cross-validation in JMP 14 to construct unified climate models to explain flowering time variation based on the whole suite of available climate factors.

#### Genotypic effects on flowering time

We first considered models in which genomic cluster was the only term included to explain patterns of variation in flowering time. We then added individual SNPs using a predictor screening approach in JMP 14, retaining SNPs that explained the highest amount of residual variance in flowering time. Finally, we used a minimum BIC model selection approach to determine which factors should be included in the final model.

#### Gene expression and flowering time

We applied a response screening approach on the complete transcriptome to identify genes whose expression patterns were most associated with flowering time using a predictor screening approach. Expression of known flowering regulators *SOC1* and *FLC* expression explained the largest amount of variation in flowering time, both at 16°C and 10°C. Many other genes with expression levels highly associated with flowering time variation were also directly associated with *SOC1* and *FLC* expression levels, and we classified genes for which *SOC1* and *FLC* expression explains >10% of their expression variance as *SOC1/FLC* associated genes. Then, after controlling for *SOC1/FLC* expression level, we identified 261 genes that explained significant variation in the flowering time residuals. After performing principal component analysis to reduce the dimensionality of their expression variation, we applied a stepwise minimum BIC model selection approach to identify which of the 10 resulting PCs were associated with flowering time variation. Finally, we identified the genes that had PC loading scores >0.20 on these components to highlight the genes that contributed most to these residual axes of flowering time variation for gene ontology analysis. We then modeled flowering time using these additional PCs alongside *FLC/SOC1* expression, and we built fully integrative models that also included genomic clusters and flowering time SNPs in JMP using a minimum BIC stepwise model selection approach. Finally, we tested for associations with climate terms between, within, and variably across clusters for *FLC/SOC1* expression, these residual gene expression PCs, and for flowering time phenotypes following the methods applied above to detect climate-GEM associations at each scale.

#### Testing flowering time models on ecologically relevant flowering time data

To test how well our models predict flowering time variation under other environmental conditions, we first calculated the predicted flowering times under 16°C and 10°C (and the difference between those values) for each ecotype based on both the genotype only model and the final model including genotypic and gene expression factors. Next, using these model predictions (rather than rerunning the model, to prevent overfitting), we tested how well these predicted flowering times in constant conditions could explain flowering time measured in growth chambers programmed to mimic the climates of natural *Arabidopsis* habitats in Spain and Sweden. We compared these models to determine 1) which constant growth chamber condition most accurately predicted flowering time in more realistic natural conditions and 2) whether gene expression (as measured at 22°C) increased the ability to predict flowering time in these more realistic conditions.

### Analysis of flowering time terms

#### Clustering

To cluster co-varying flowering-time associated SNPs (from ref. 61) as well as flowering time gene expression and climate terms of interest, we created a similarity matrix between terms based on the absolute value of the correlation between terms across all individuals and applied Ward-based clustering to this matrix with the R package pvclust (92). To confirm clusters of closely linked terms, we also performed density-based clustering with eps=1.2 in dbscan (93, 94).

#### Structural equation modeling

We performed pathway analysis with the structural equation modeling software Ωnyx (95) to assess the magnitude and direction of intermediary causal relationships impacting the observed associations between climate terms, genomic cluster and flowering time SNPs, gene expression, and flowering time. We removed associations between levels with effect sizes < 0.02.

## Acknowledgments

The authors thank Emily Josephs and Tom Juenger for helpful conversations during the development of this work, and we thank the Blackman lab and David Des Marais for their generous feedback on early versions of this manuscript. Support for this project was provided by the University of California, Berkeley, National Science Foundation grants to BKB (IOS-1558035, DEB-1640788), and a National Institutes of Health Ruth L. Kirschstein National Research Service Award Individual Postdoctoral Fellowship to JMC (F32 GM125244-01). The authors thank the *Arabidopsis* community for the availability and accessibility of transcriptomic, genomic, phenotypic and geospatial information for all accessions.

## Author Contributions

JMC conceived the project and analytical framework. JMC, MA, and BKB generated hypotheses and identified tools to test these questions. JMC and MA performed the data analyses. JMC, MA, and BKB wrote the manuscript.

## Supplementary Information for

**Fig. S1.**
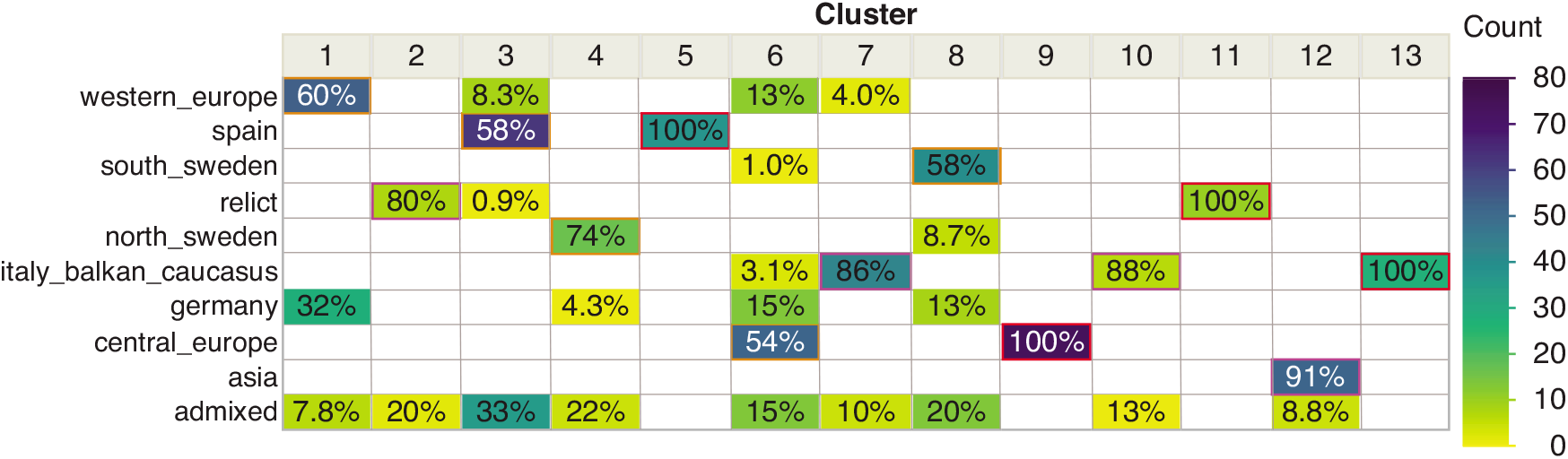
Congruence of 13 genomic clusters identified and used in this study with the ancestry groups found by the 1001 Genomes Consortium [39]. Percentages indicate the percent of a given genomic cluster in each ancestry group, color coded by the number of accessions in a given group.

**Fig. S2.**
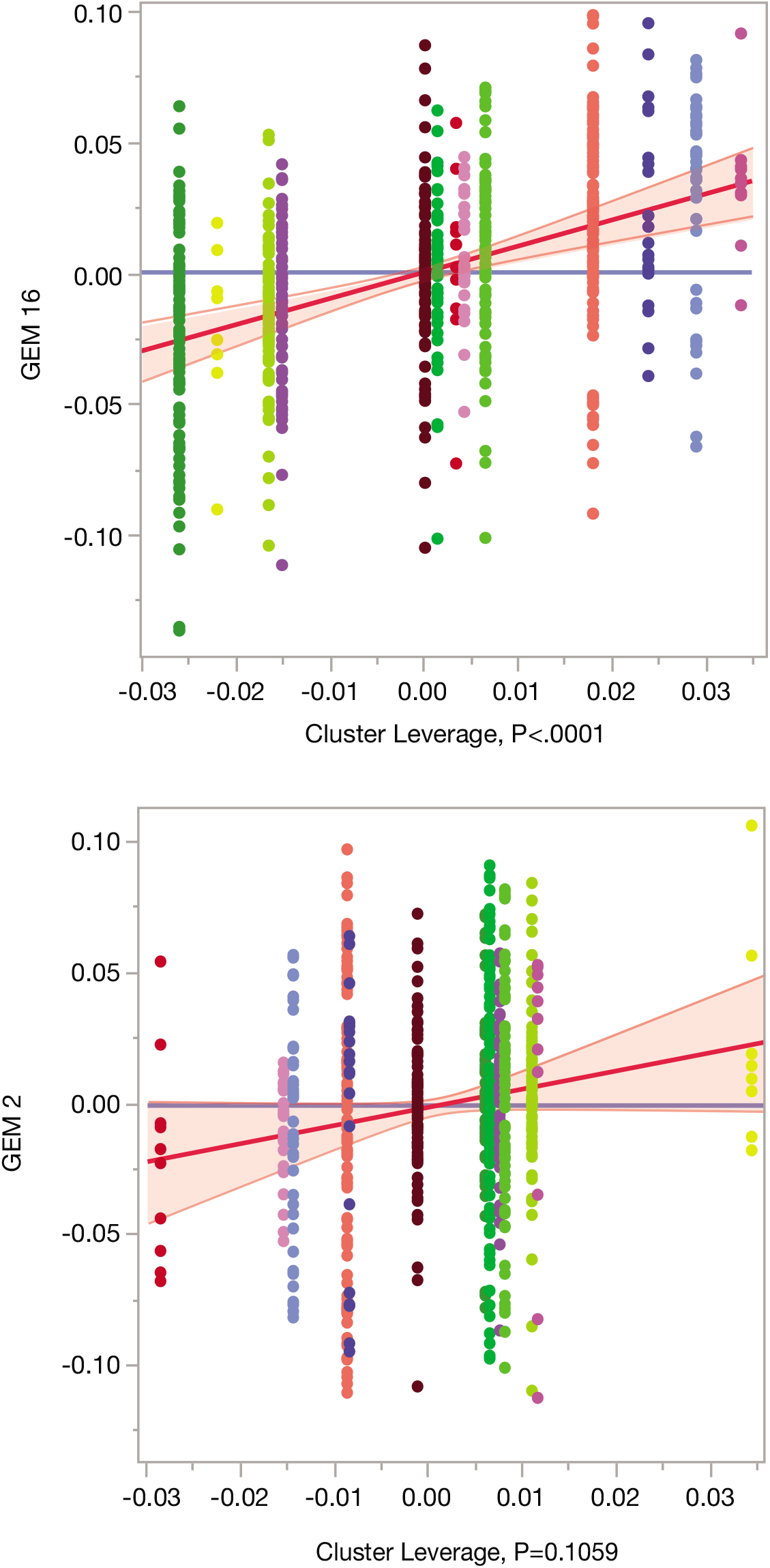
Least squares regression plots of GEM16 and GEM2 eigengene expression values on genomic cluster. Points colored by genomic cluster. Genomic cluster explains 20% of the variation for GEM16 (p<0.0001), but only 2.8% of the variance for GEM2 (p=0.11).

**Figure S3.**
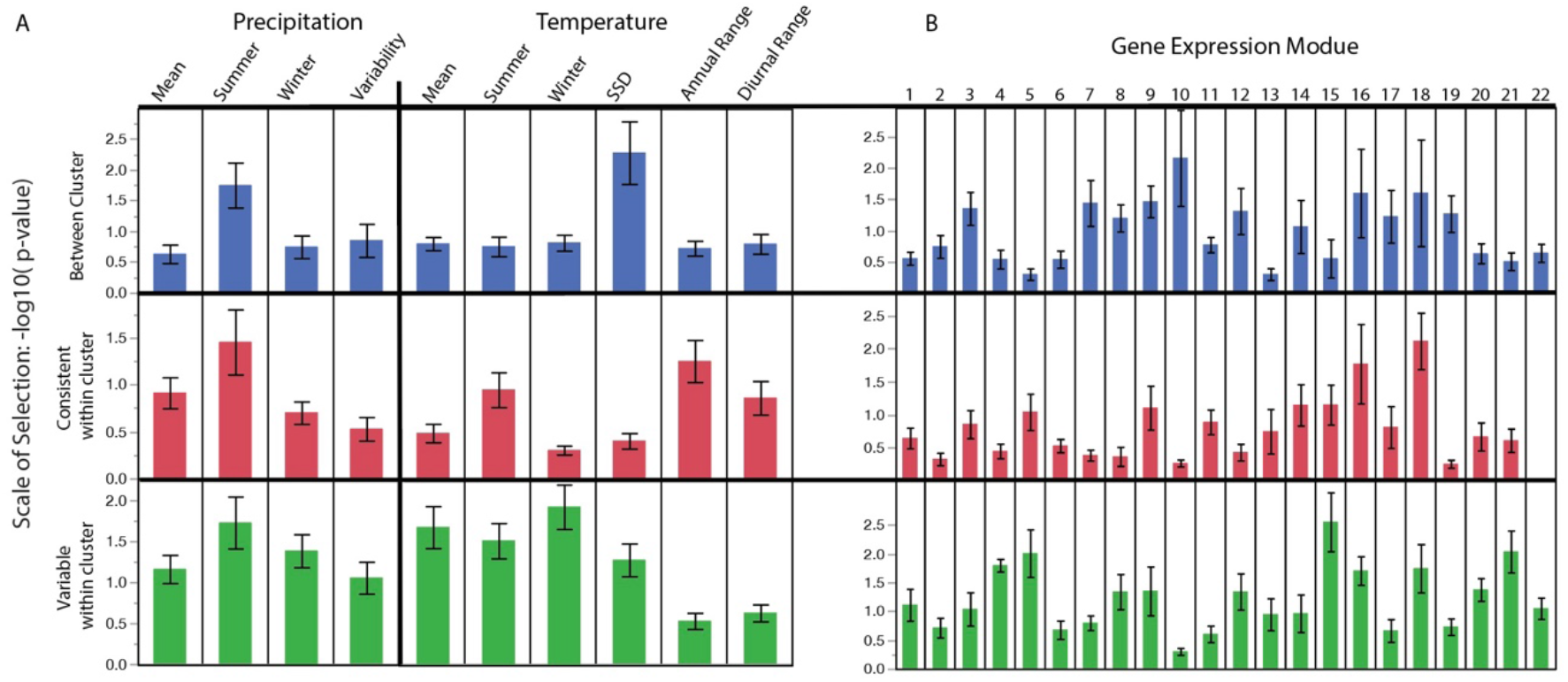
Means of the negative log significance for GEM ~ Climate associations across three scales grouped by (A) climate term and (B) gene expression module.

**Figure S4.**
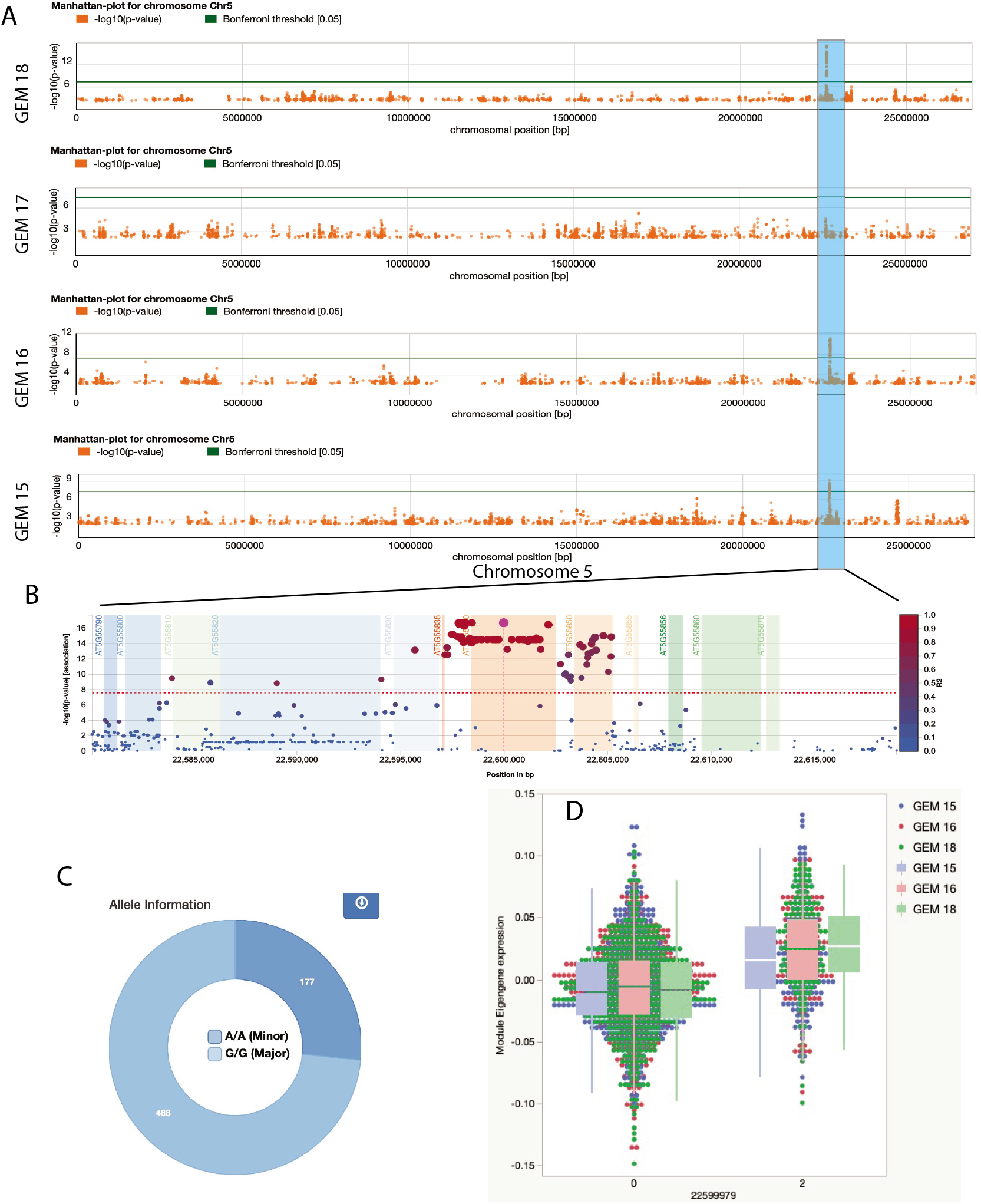
Module expression GWAS finds a major eQTL explaining variation in the expression of GEMs 15, 16, and 18. (A) LOD significance plots show overlapping genome wide significant peak for GEMs 15,16, 18, but not 17. (B) Fine-scale plot of SNP associations reveals peak is primarily delimited to an ~8kb region. Regions spanned by annotations in this interval are indicated by shaded background blocks. (C) For the top SNP in this region (5_22599979) 468 accessions have the reference G/G allele, and 177 have the alt A/A allele. (D) Individuals homozygous for the alt A/A allele (2) have significantly higher expression of modules 15, 16, and 18.

**Figure S5.**
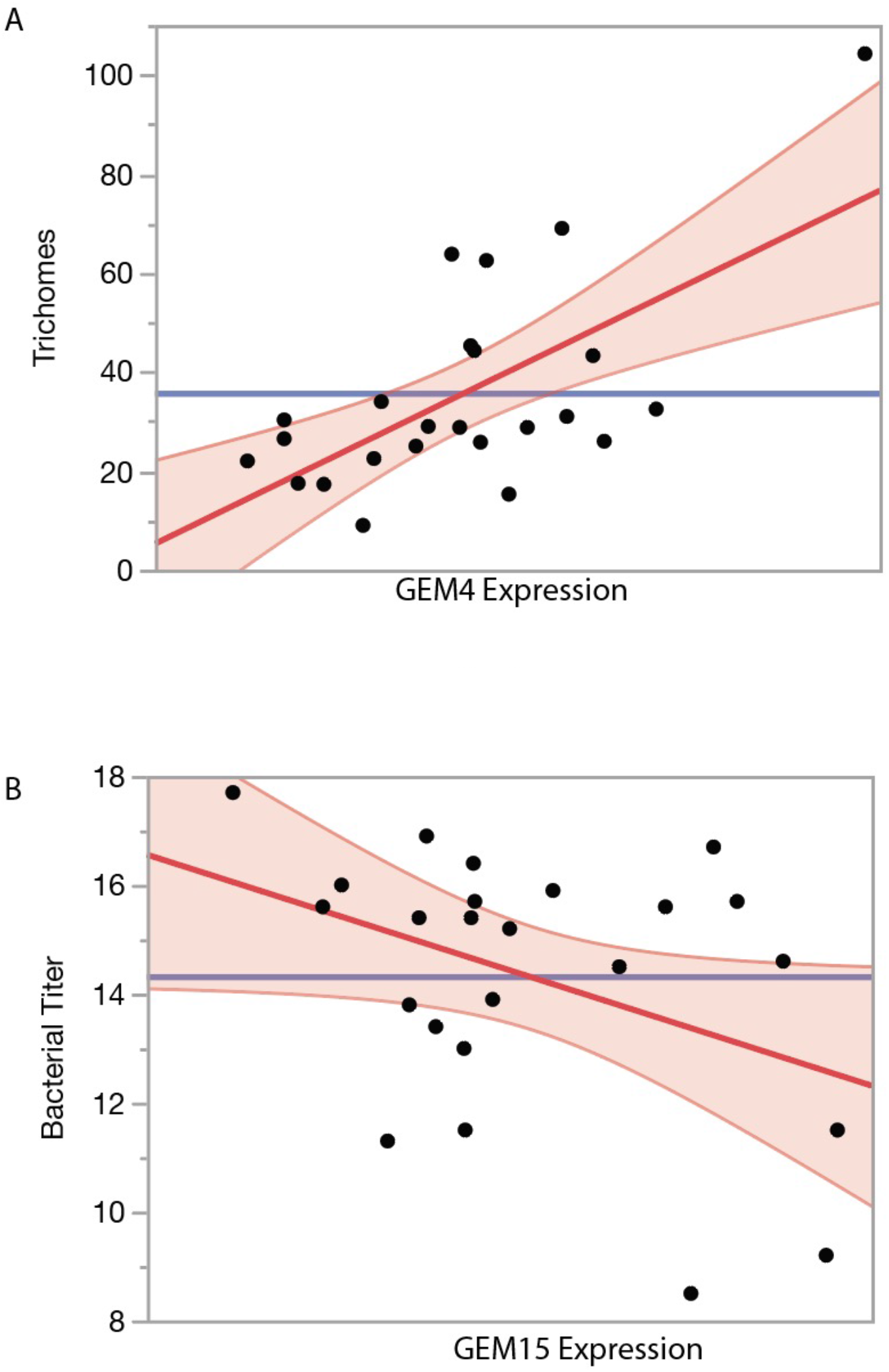
Standard least squares regression of (A) accession trichomes on GEM4 (p=0.0006, R^2^=0.42) and (B) bacterial titer on GEM15 (p=0.05. R^2^=0.16).

**Figure S6.**
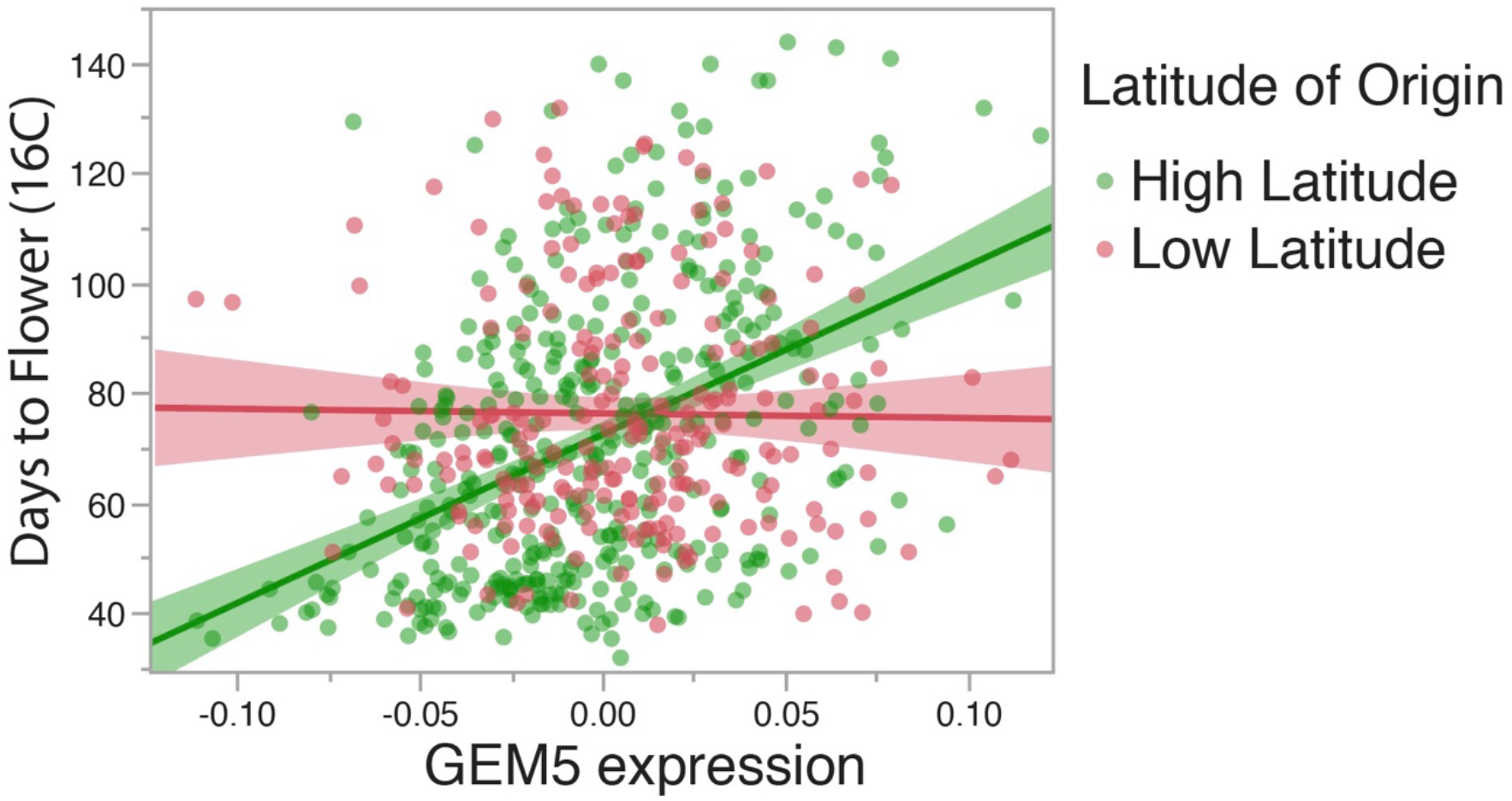
Associations between eigengene expression values GEM5 (enriched for defense-related genes) and FT16 are highly dependent on accession latitude of origin. High latitude lines (above 45°N) have a positive correlation between GEM5 expression and FT16 (R^2^=0.25, p<0.0001); lines from lower latitudes show no significant association between these two variables (R^2^=0.01, p=0.22).

**Figure S7.**
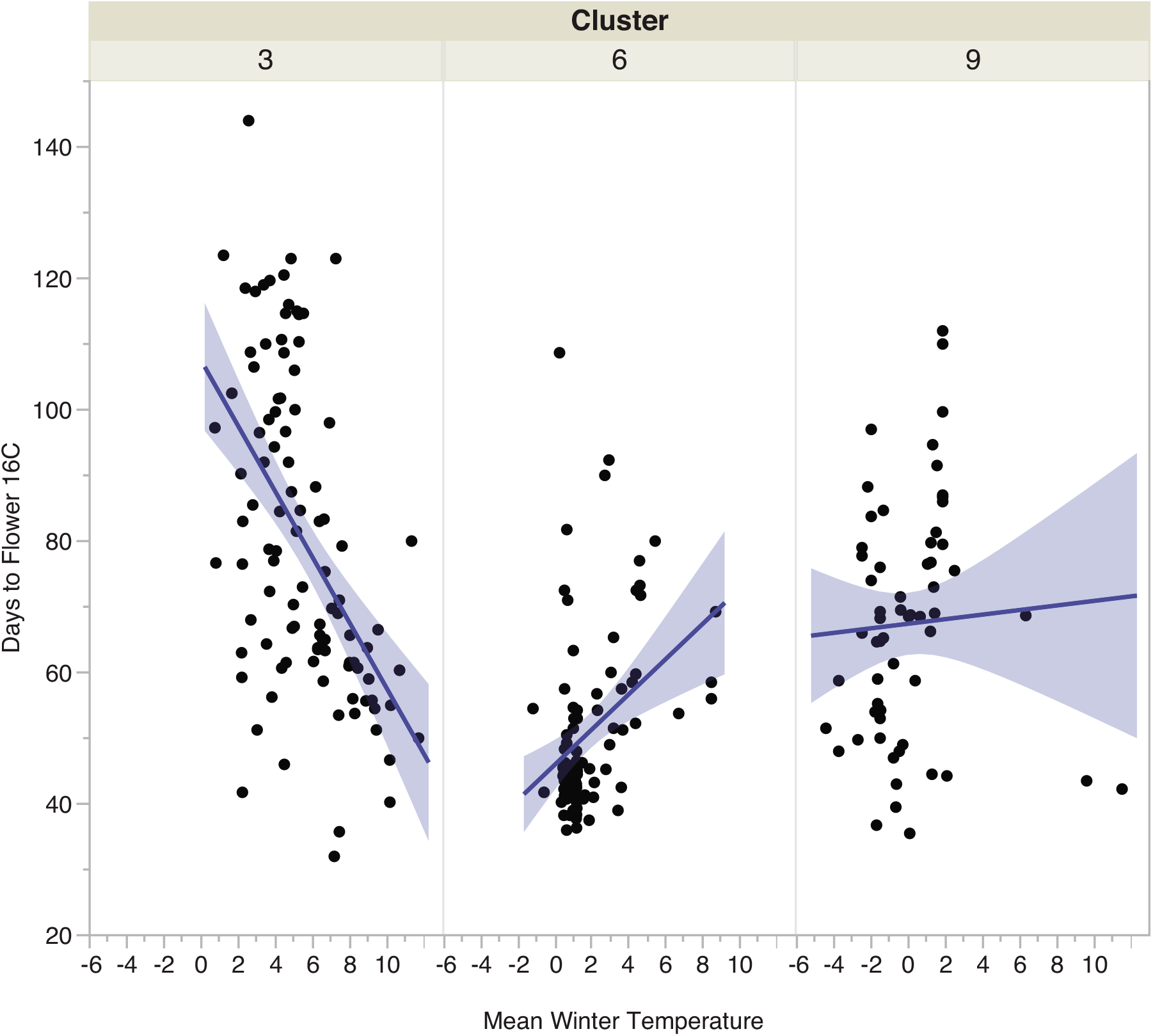
Example of the variability in cluster-specific associations of flowering time with climate parameters. Associations between mean winter temperature and FT16 are highly dependent upon the specific genomic group that on considers. There is a highly significant (p<0.0001, R^2^=0.26) negative association with mean winter temperature and days to flower for cluster 3, a moderately significant association within cluster 6 (p=0.002, R^2^=0.11), and no evidence of an association within cluster 9 (p=0.6, R^2^=0.003).

**Figure S8.**
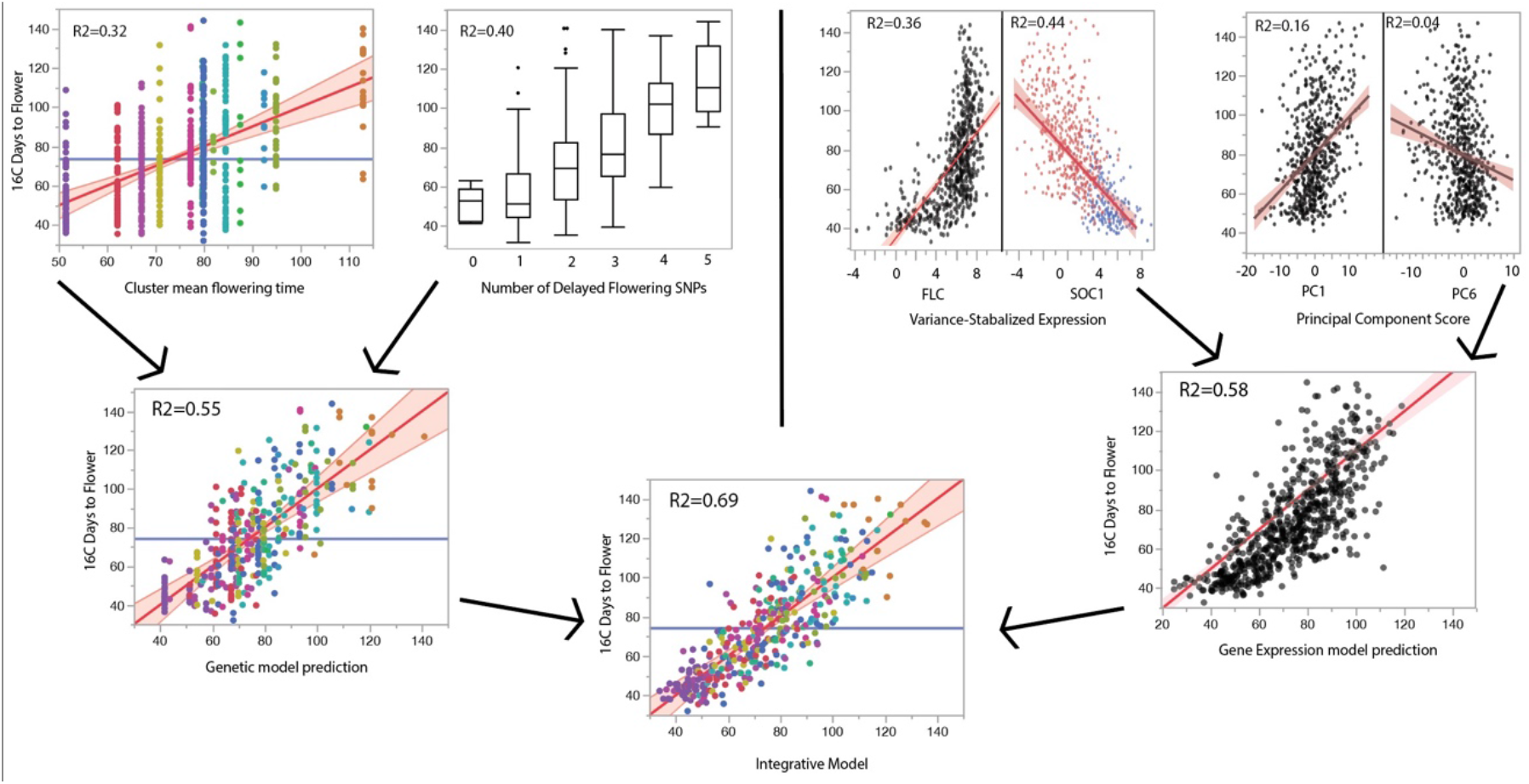
Schematic illustrating how various data sources were combined in the development of the integrative flowering time model. On the left, genomic cluster and individual genotype for flowering time SNPs each explain significant portions of flowering time variation, and explain a greater portion of variance when modeled together. Colors represent different genomic clusters. On the right, models showing the relationship between gene expression and flowering time variation including top terms *FLC* and *SOC1* as well as the expression components associated with flowering time residuals after controlling for *FLC/SOC1* expression, PC1 and PC6. Combining various classes of data increases model capacity to explain flowering time variation.

**Figure S9.**
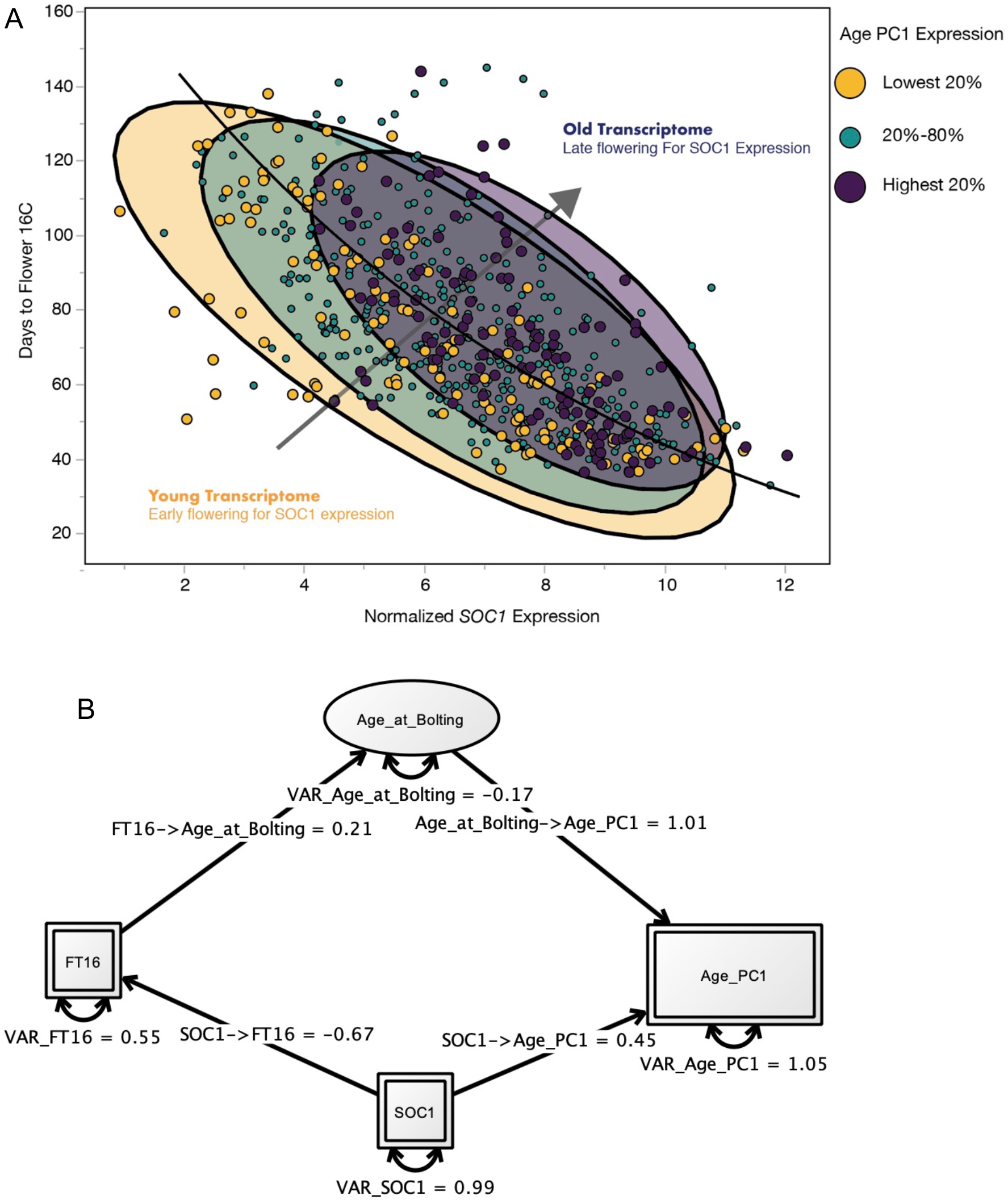
Relationship between *SOC1* expression and days to flower can be further partitioned based on the expression of aging associated genes. Accessions with higher expression of age-associated PC1 tend to flower later than would be expected based upon their *SOC1* expression alone (tend to be located above the curve of fit). The opposite pattern is seen for individuals with low expression of age PC1. (B) Structural equation model with *SOC1* expression impacting both FT16 as well as age PC1.

**Figure S10.**
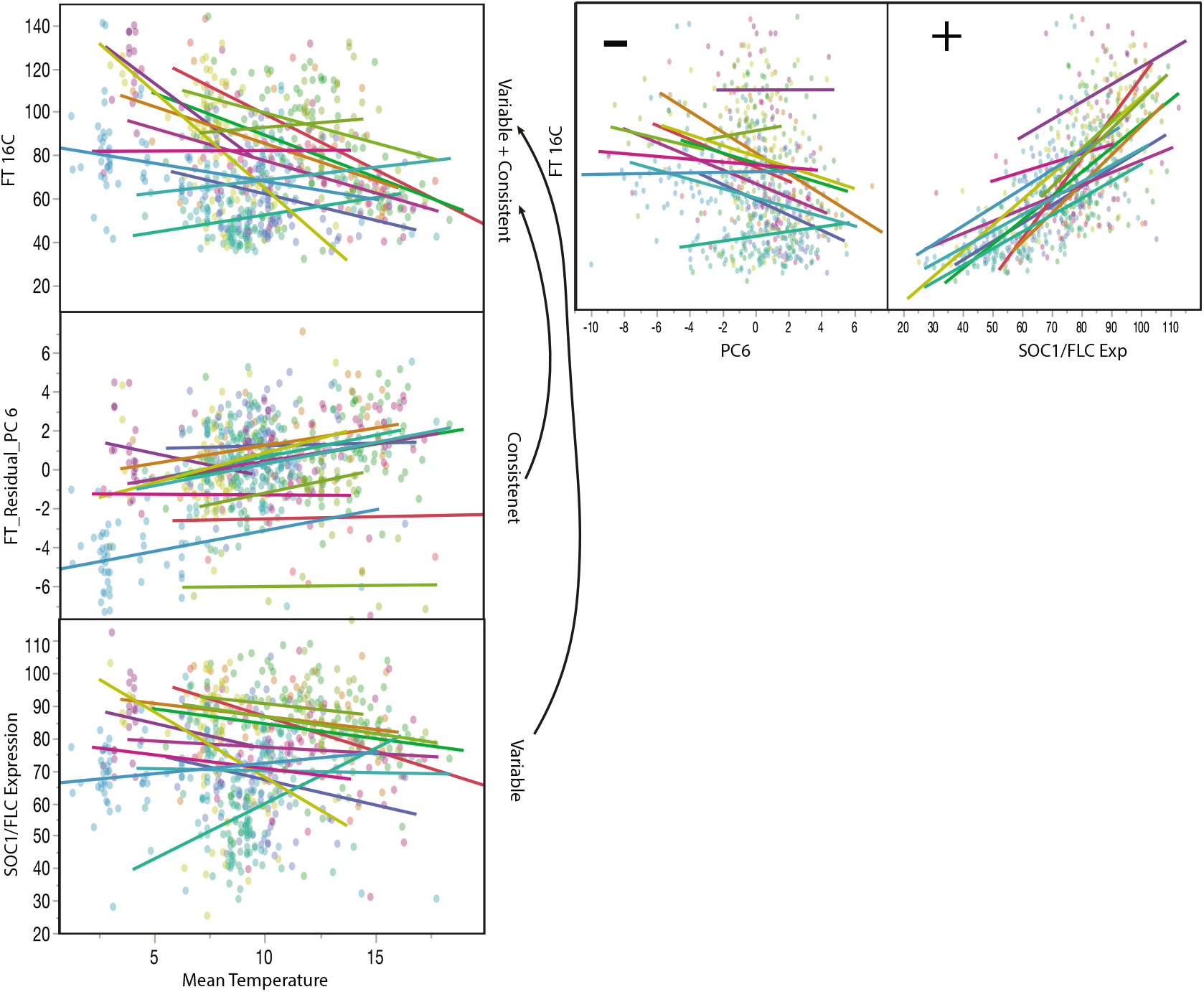
Mean annual temperature shows evidence of both consistent and genomic cluster dependent associations with flowering time. There is a largely consistent positive association between mean annual temperature and expression PC6, and a largely consistent negative association between this term and flowering time in 16C, driving a subtle but consistent negative association between mean annual temperature and flowering time. In contrast, the association between mean annual temperature and *FLC* and *SOC1* expression (here standardized to expected days to flower for a given *FLC/SOC1* level) is highly variable among genomic clusters (shown in different colors). This term is highly positively associated with days to flower, in turn leading to highly significant cluster dependent variable associations between mean temperature and flowering time. These two expression terms combine to give rise to detectable evidence of a consistent negative association between mean annual and temperature and flowering time (driven by PC6), but also a large amount of between genomic cluster variation for these patterns (driven by *FLC/SOC1).*

**Figure S11.**
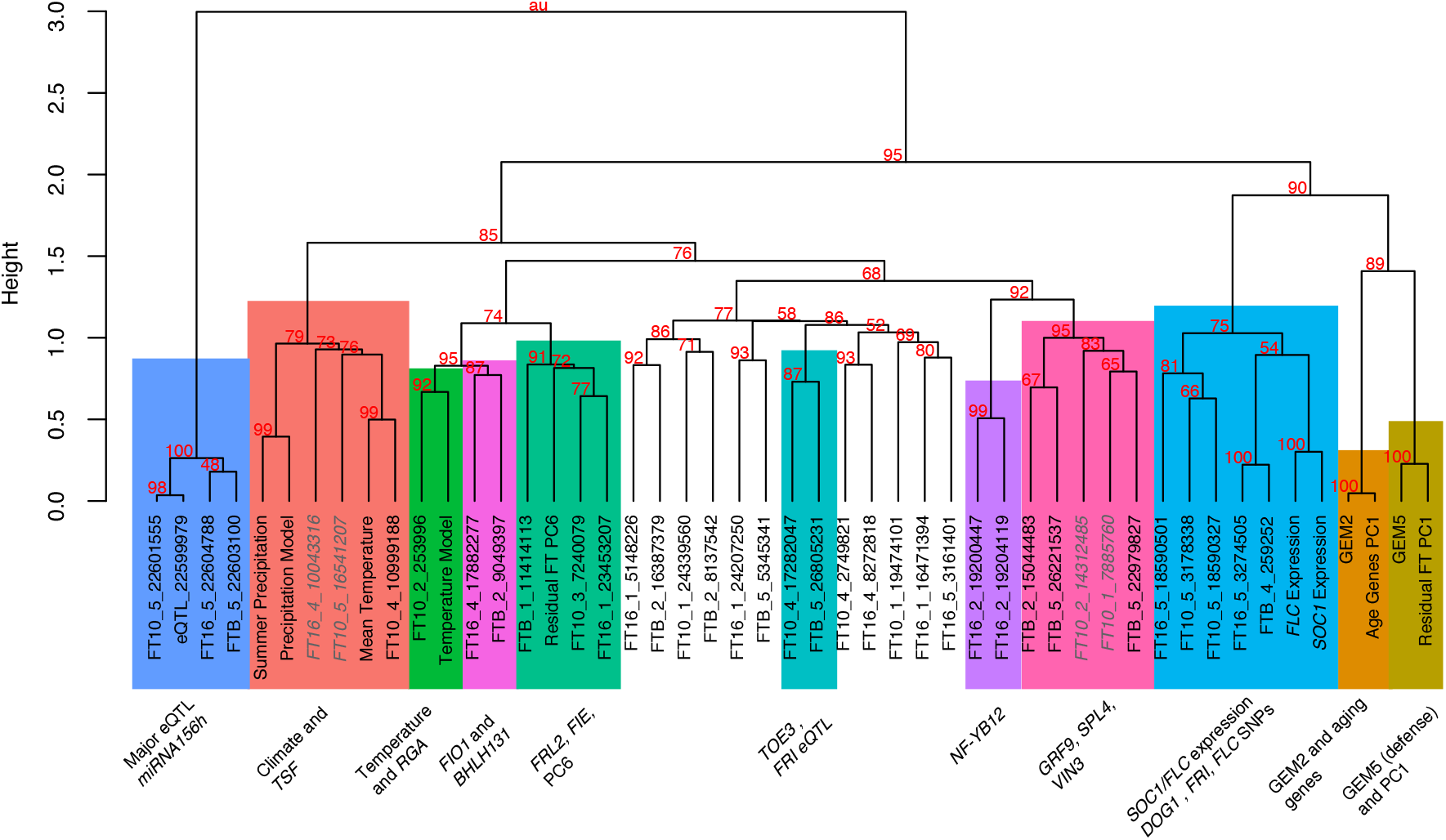
Ward based clustering of flowering time associated terms. R pvclust used to calculate bootstrap confidence, and Rdbscan to identify significant clusters (colored bins).

## Notes

http://signal.salk.edu/1001.php

https://1001genomes.org/data-center.html

### Competing Interest Statement

The authors have declared no competing interest.

## References

1. J. Huxley, Clines: an Auxiliary Taxonomic Principle. Nature 142, 219–220 (1938).

2. J. A. Endler, Gene flow and population differentiation: studies of clines suggest that differentiation along environmental gradients may be independent of gene flow. Science 179, 243–250 (1973).

3. T. Dobzhansky, Genetics of natural populations; experiments on chromosomes of Drosophila pseudoobscura from different geographic regions. Genetics 33, 588–602 (1948).

4. A. R. Place, D. A. Powers, Genetic variation and relative catalytic efficiencies: lactate dehydrogenase B allozymes of *Fundulus heteroclitus*. Proc. Natl. Acad. Sci. U. S. A. 76, 2354–2358 (1979).

5. R. G. Collevatti, et al., A genome-wide scan shows evidence for local adaptation in a widespread keystone Neotropical forest tree. Heredity 123, 117–137 (2019).

6. J. K. Pritchard, J. K. Pickrell, G. Coop, The genetics of human adaptation: hard sweeps, soft sweeps, and polygenic adaptation. Curr. Biol. 20, R208–15 (2010).

7. M. Takou, et al., Linking genes with ecological strategies in *Arabidopsis thaliana*. J. Exp. Bot. 70, 1141–1151 (2019).

8. N. J. Kooyers, A. B. Greenlee, J. M. Colicchio, M. Oh, B. K. Blackman, Replicate altitudinal clines reveal that evolutionary flexibility underlies adaptation to drought stress in annual *Mimulus guttatus*. New Phytol. 206, 152–165 (2015).

9. M. Wellenreuther, B. Hansson, Detecting Polygenic Evolution: Problems, Pitfalls, and Promises. Trends Genet. 32, 155–164 (2016).

10. C. O’Brien, W. E. Bradshaw, C. M. Holzapfel, Testing for causality in covarying traits: genes and latitude in a molecular world. Mol. Ecol. 20, 2471–6 (2011).

11. B. Brachi, et al., Investigation of the geographical scale of adaptive phenological variation and its underlying genetics in *Arabidopsis thaliana*. Mol. Ecol. 22, 4222–4240 (2013).

12. D. Tabas-Madrid, B. Méndez-Vigo, Genome-wide signatures of flowering adaptation to climate temperature: Regional analyses in a highly diverse native range of *Arabidopsis thaliana*. Plant Cell Environ. (2018).

13. C. Oberprieler, C. Zimmer, M. Bog, Are there morphological and life-history traits under climate-dependent differential selection in S Tunesian Diplotaxis harra (Forssk.) Boiss.(Brassicaceae) populations? Ecol. Evol. 8, 1047–1062 (2018).

14. A. A. Fikas, B. P. Dilkes, I. Baxter, Multivariate analysis reveals environmental and genetic determinants of element covariation in the maize grain ionome. Plant Direct 3, e00139 (2019).

15. N. J. Kooyers, K. M. Olsen, Adaptive cyanogenesis clines evolve recurrently through geographical sorting of existing gene deletions. J. Evol. Biol. 27, 2554–2558 (2014).

16. T. Kawakatsu, et al., Epigenomic Diversity in a Global Collection of *Arabidopsis thaliana* Accessions. Cell 166, 492–505 (2016).

17. H. B. Fraser, A. M. Moses, E. E. Schadt, Evidence for widespread adaptive evolution of gene expression in budding yeast. Proc. Natl. Acad. Sci. U. S. A. 107, 2977–2982 (2010).

18. A. Whitehead, D. L. Crawford, Neutral and adaptive variation in gene expression. Proc. Natl. Acad. Sci. U. S. A. 103, 5425–5430 (2006).

19. K. L. Mack, M. A. Ballinger, M. Phifer-Rixey, M. W. Nachman, Gene regulation underlies environmental adaptation in house mice. Genome Res. 28, 1636–1645 (2018).

20. S. C. Groen, et al., The strength and pattern of natural selection on gene expression in rice. Nature 578, 572–576 (2020).

21. D. Porcelli, et al., Gene expression clines reveal local adaptation and associated trade-offs at a continental scale. Sci. Rep. 6, 32975 (2016).

22. S. A. Signor, S. V. Nuzhdin, The Evolution of Gene Expression in *cis* and *trans*. Trends Genet. 34, 532–544 (2018).

23. I. G. Romero, I. Ruvinsky, Y. Gilad, Comparative studies of gene expression and the evolution of gene regulation. Nat. Rev. Genet. 13, 505–516 (2012).

24. J. H. Bullard, Y. Mostovoy, S. Dudoit, R. B. Brem, Polygenic and directional regulatory evolution across pathways in *Saccharomyces*. Proc. Natl. Acad. Sci. U. S. A. 107, 5058–5063 (2010).

25. M. A. House, C. K. Griswold, L. N. Lukens, Evidence for selection on gene expression in cultivated rice *(Oryza sativa)*. Mol. Biol. Evol. 31, 1514–1525 (2014).

26. H. B. Fraser, et al., Systematic detection of polygenic *cis*-regulatory evolution. PLoS Genet. 7, e1002023 (2011).

27. P. A. Lavin, J. D. Mcphail, Parapatric lake and stream sticklebacks on northern Vancouver Island: disjunct distribution or parallel evolution? Can. J. Zool. 71, 11–17 (1993).

28. P. Ralph, G. Coop, Parallel adaptation: one or many waves of advance of an advantageous allele? Genetics 186, 647–668 (2010).

29. S. Hoban, et al., Finding the Genomic Basis of Local Adaptation: Pitfalls, Practical Solutions, and Future Directions. Am. Nat. 188, 379–397 (2016).

30. B. K. Blackman, S. D. Michaels, L. H. Rieseberg, Connecting the sun to flowering in sunflower adaptation. Mol. Ecol. 20, 3503–3512 (2011).

31. L. Zhao, J. Wit, N. Svetec, D. J. Begun, Parallel Gene Expression Differences between Low and High Latitude Populations of *Drosophila melanogaster* and *D. simulans*. PLoS Genet. 11, e1005184 (2015).

32. L. U. Gleason, R. S. Burton, RNA-seq reveals regional differences in transcriptome response to heat stress in the marine snail *Chlorostoma funebralis*. Mol. Ecol. 24, 610–627 (2015).

33. D. L. Des Marais, R. F. Guerrero, J. R. Lasky, S. V. Scarpino, Topological features of a gene co-expression network predict patterns of natural diversity in environmental response. Proc. Biol. Sci. 284 (2017).

34. S. C. Campbell-Staton, et al., Parallel selection on thermal physiology facilitates repeated adaptation of city lizards to urban heat islands. Nat. Ecol. Evol. 4, 652–658 (2020).

35. M. Akman, J. E. Carlson, K. E. Holsinger, A. M. Latimer, Transcriptome sequencing reveals population differentiation in gene expression linked to functional traits and environmental gradients in the South African shrub *Protea repens*. New Phytol. 210, 295–309 (2016).

36. M. Akman, J. E. Carlson, A. M. Latimer, Climate gradients explain population-level divergence in drought-induced plasticity of functional traits and gene expression in a South African *Protea*. Mol. Ecol. (2020) https:/doi.org/10.1111/mec.15705.

37. M. A. Kost, et al., Differentiated transcriptional signatures in the maize landraces of Chiapas, Mexico. BMC Genomics 18, 707 (2017).

38. M. Zhu, et al., WGCNA Analysis of Salt-Responsive Core Transcriptome Identifies Novel Hub Genes in Rice. Genes 10 (2019).

39. Q. Wang, et al., Identification of key genes and modules in response to cadmium stress in different rice varieties and stem nodes by weighted gene co-expression network analysis. Sci. Rep. 10, 9525 (2020).

40. M. A. Kost, et al., Transcriptional differentiation of UV-B protectant genes in maize landraces spanning an elevational gradient in Chiapas, Mexico. Evol. Appl. 13, 1949–1967 (2020).

41. E. Sasaki, F. Frommlet, M. Nordborg, GWAS with Heterogeneous Data: Estimating the Fraction of Phenotypic Variation Mediated by Gene Expression Data. G3 8, 3059–3068 (2018).

42. Y. Zan, Ö. Carlborg, A multilocus association analysis method integrating phenotype and expression data reveals multiple novel associations to flowering time variation in wild-collected *Arabidopsis thaliana*. Mol. Ecol. Resour. 18, 798–808 (2018).

43. 1001 Genomes Consortium, 1,135 Genomes Reveal the Global Pattern of Polymorphism in *Arabidopsis thaliana*. Cell 166, 481–491 (2016).

44. P. Langfelder, S. Horvath, WGCNA: an R package for weighted correlation network analysis. BMC Bioinformatics 9, 559 (2008).

45. G. Wu, R. S. Poethig, Temporal regulation of shoot development in *Arabidopsis thaliana* by miR156 and its target SPL3. Development 133, 3539–3547 (2006).

46. N. Mähler, et al., Gene co-expression network connectivity is an important determinant of selective constraint. PLoS Genet. 13, e1006402 (2017).

47. T. E. Scarpeci, V. S. Frea, M. I. Zanor, E. M. Valle, Overexpression of AtERF019 delays plant growth and senescence, and improves drought tolerance in *Arabidopsis*. J. Exp. Bot. 68, 673–685 (2017).

48. M. Salehin, et al., Auxin-sensitive Aux/IAA proteins mediate drought tolerance in *Arabidopsis* by regulating glucosinolate levels. Nat. Commun. 10, 4021 (2019).

49. Y. Ding, et al., Four distinct types of dehydration stress memory genes in *Arabidopsis thaliana*. BMC Plant Biol. 13, 229 (2013).

50. D. Huang, W. Wu, S. R. Abrams, A. J. Cutler, The relationship of drought-related gene expression in *Arabidopsis thaliana* to hormonal and environmental factors. J. Exp. Bot. 59, 2991–3007 (2008).

51. S. Atwell, et al., Genome-wide association study of 107 phenotypes in *Arabidopsis thaliana* inbred lines. Nature 465, 627–631 (2010).

52. B. Brachi, et al., Coselected genes determine adaptive variation in herbivore resistance throughout the native range of *Arabidopsis thaliana*. Proc. Natl. Acad. Sci. U. S. A. 112, 4032–4037 (2015).

53. A. M. Wentzell, et al., Linking metabolic QTLs with network and *cis*-eQTLs controlling biosynthetic pathways. PLoS Genet. 3, 1687–1701 (2007).

54. S. Glander, et al., Assortment of Flowering Time and Immunity Alleles in Natural *Arabidopsis thaliana* Populations Suggests Immunity and Vegetative Lifespan Strategies Coevolve. Genome Biol. Evol. 10, 2278–2291 (2018).

55. R. Lyons, et al., Investigating the Association between Flowering Time and Defense in the *Arabidopsis thaliana-Fusarium oxysporum* Interaction. PLoS One 10, e0127699 (2015).

56. B. Li, et al., Epistatic Transcription Factor Networks Differentially Modulate *Arabidopsis* Growth and Defense. Genetics 214, 529–541 (2020).

57. J. G. Monroe, et al., Drought adaptation in *Arabidopsis thaliana* by extensive genetic loss-of-function. Elife 7, e41038 (2018).

58. J. R. Stinchcombe, et al., A latitudinal cline in flowering time in *Arabidopsis thaliana* modulated by the flowering time gene *FRIGIDA*. Proc. Natl. Acad. Sci. U. S. A. 101, 4712–4717 (2004).

59. C. Shindo, et al., Role of *FRIGIDA* and *FLOWERING LOCUS C* in determining variation in flowering time of *Arabidopsis*. Plant Physiol. 138, 1163–1173 (2005).

60. M. Exposito-Alonso, A. C. Brennan, C. Alonso-Blanco, F. X. Picó, Spatio-temporal variation in fitness responses to contrasting environments in *Arabidopsis thaliana:* fitness responses to novel environments. Evolution 71, 550 (2018).

61. Y. Zan, Ö. Carlborg, A Polygenic Genetic Architecture of Flowering Time in the Worldwide *Arabidopsis thaliana* Population. Mol. Biol. Evol. 36, 141–154 (2019).

62. W. Deng, et al., FLOWERING LOCUS C (FLC) regulates development pathways throughout the life cycle of *Arabidopsis*. Proc. Natl. Acad. Sci. U. S. A. 108, 6680–6685 (2011).

63. S. K. Yoo, et al., *CONSTANS* activates *SUPPRESSOR OF OVEREXPRESSION OF CONSTANS 1* through *FLOWERING LOCUS T* to promote flowering in *Arabidopsis*. Plant Physiol. 139, 770–778 (2005).

64. P. Y. Hsu, S. L. Harmer, Wheels within wheels: the plant circadian system. Trends Plant Sci. 19, 240–249 (2014).

65. Y. Toda, T. Kudo, T. Kinoshita, N. Nakamichi, Evolutionary Insight into the Clock-Associated PRR5 Transcriptional Network of Flowering Plants. Sci. Rep. 9, 2983 (2019).

66. N. Nakamichi, et al., Transcriptional repressor PRR5 directly regulates clock-output pathways. Proc. Natl. Acad. Sci. U. S. A. 109, 17123–17128 (2012).

67. F. Fornara, et al., *Arabidopsis* DOF transcription factors act redundantly to reduce *CONSTANS* expression and are essential for a photoperiodic flowering response. Dev. Cell 17, 75–86 (2009).

68. J. Lee, I. Lee, Regulation and function of SOC1, a flowering pathway integrator. J. Exp. Bot. 61, 2247–2254 (2010).

69. T. Imaizumi, H. G. Tran, T. E. Swartz, W. R. Briggs, S. A. Kay, FKF1 is essential for photoperiodic-specific light signalling in *Arabidopsis*. Nature 426, 302–306 (2003).

70. S. R. Hepworth, F. Valverde, D. Ravenscroft, A. Mouradov, G. Coupland, Antagonistic regulation of flowering-time gene *SOC1* by CONSTANS and FLC via separate promoter motifs. EMBO J. 21, 4327–4337 (2002).

71. S. Melzer, et al., Flowering-time genes modulate meristem determinacy and growth form in *Arabidopsis thaliana*. Nat. Genet. 40, 1489–1492 (2008).

72. J. Li, G. Brader, E. T. Palva, The WRKY70 transcription factor: a node of convergence for jasmonate-mediated and salicylate-mediated signals in plant defense. Plant Cell 16, 319–331 (2004).

73. A. Martínez-Berdeja, et al., Functional variants of *DOG1* control seed chilling responses and variation in seasonal life-history strategies in *Arabidopsis thaliana*. Proc. Natl. Acad. Sci. U. S. A. 117, 2526–2534 (2020).

74. H. Huo, S. Wei, K. J. Bradford, *DELAY OF GERMINATION1 (DOG1)* regulates both seed dormancy and flowering time through microRNA pathways. Proc. Natl. Acad. Sci. U. S. A. 113, E2199–E2206 (2016).

75. J. Chen, et al., *Arabidopsis* WRKY46, WRKY54, and WRKY70 Transcription Factors Are Involved in Brassinosteroid-Regulated Plant Growth and Drought Responses. Plant Cell 29, 1425–1439 (2017).

76. B. Ulker, M. Shahid Mukhtar, I. E. Somssich, The WRKY70 transcription factor of *Arabidopsis* influences both the plant senescence and defense signaling pathways. Planta 226, 125–137 (2007).

77. R. B. Deal, C. N. Topp, E. C. McKinney, R. B. Meagher, Repression of flowering in *Arabidopsis* requires activation of *FLOWERING LOCUS C* expression by the histone variant H2A.Z. Plant Cell 19, 74–83 (2007).

78. S. V. Kumar, et al., Transcription factor PIF4 controls the thermosensory activation of flowering. Nature 484, 242–245 (2012).

79. A. C. Velásquez, C. D. M. Castroverde, S. Y. He, Plant–Pathogen Warfare under Changing Climate Conditions. Curr. Biol. 28, R619–R634 (2018).

80. T. Srikant, A. Wibowo, R. Schwab, D. Weigel, Position-dependent effects of cytosine methylation on FWA expression in *Arabidopsis thaliana*. BioRxiv, 774281 (2019).

81. M. D. Robinson, D. J. McCarthy, G. K. Smyth, edgeR: a Bioconductor package for differential expression analysis of digital gene expression data. Bioinformatics 26, 139–140 (2010).

82. M. E. Ritchie, et al., limma powers differential expression analyses for RNA-sequencing and microarray studies. Nucleic Acids Res. 43, e47 (2015).

83. X. Zheng, et al., A high-performance computing toolset for relatedness and principal component analysis of SNP data. Bioinformatics 28, 3326–3328 (2012).

84. SAS Institute Inc., Cary, NC, JMP®, Version 14.0 (1989-2019).

85. A. M. Hancock, et al., Adaptation to climate across the *Arabidopsis thaliana* genome. Science 334, 83–86 (2011).

86. M. Exposito-Alonso, et al., Genomic basis and evolutionary potential for extreme drought adaptation in *Arabidopsis thaliana*. Nat. Ecol. Evol. 2, 352–358 (2018).

87. Y. Li, P. Roycewicz, E. Smith, J. O. Borevitz, Genetics of local adaptation in the laboratory: flowering time quantitative trait loci under geographic and seasonal conditions in *Arabidopsis*. PLoS One 1, e105 (2006).

88. S. E. Fick, R. J. Hijmans, WorldClim 2: new 1-km spatial resolution climate surfaces for global land areas. Int. J. Climatol. 37, 4302–4315 (2017).

89. J. Jin, et al., PlantTFDB 4.0: toward a central hub for transcription factors and regulatory interactions in plants. Nucleic Acids Res. 45, D1040–D1045 (2017).

90. P. D. Thomas, et al., PANTHER: a library of protein families and subfamilies indexed by function. Genome Res. 13, 2129–2141 (2003).

91. D. G. Grimm, et al., easyGWAS: A Cloud-Based Platform for Comparing the Results of Genome-Wide Association Studies. Plant Cell 29, 5–19 (2017).

92. R. Suzuki, H. Shimodaira, Pvclust: an R package for assessing the uncertainty in hierarchical clustering. Bioinformatics 22, 1540–1542 (2006).

93. M. Ester, H.-P. Kriegel, J. Sander, X. Xu, A density-based algorithm for discovering clusters in large spatial databases with noise in KDD’96: Proceedings of the Second International Conference on Knowledge Discovery and Data Mining, (aaai.org, 1996), pp. 226–231.

94. M. Hahsler, M. Piekenbrock, S. Arya, D. Mount, dbscan: Density Based Clustering of Applications with Noise (DBSCAN) and Related Algorithms. R package version, 1–0 (2017).

95. T. von Oertzen, A. M. Brandmaier, S. Tsang, Structural equation modeling with Ωnyx. Struct. Equ. Modeling 22, 148–161 (2015).

